# Smart Distributed Data Factory: Volunteer Computing Platform for Active Learning-Driven Molecular Data Acquisition

**DOI:** 10.1101/2024.10.22.619651

**Authors:** Tsolak Ghukasyan, Vahagn Altunyan, Aram Bughdaryan, Tigran Aghajanyan, Khachik Smbatyan, Garegin A. Papoian, Garik Petrosyan

## Abstract

This paper presents the Smart Distributed Data Factory (SDDF), an AI-driven distributed computing platform designed to address challenges in drug discovery by creating comprehensive datasets of molecular conformations and their properties. SDDF uses volunteer computing, leveraging the processing power of personal computers worldwide to accelerate quantum chemistry (DFT) calculations. To tackle the vast chemical space and limited high-quality data, SDDF employs an ensemble of machine learning models to predict molecular properties and selectively choose the most challenging data points for further DFT calculations. The platform also generates new molecular conformations using molecular dynamics with the forces derived from these models. SDDF makes several contributions: the volunteer computing platform for DFT calculations; an active learning framework for constructing a dataset of molecular conformations; a large public dataset of diverse ENAMINE molecules with calculated energies; an ensemble of state-of-the-art ML models for accurate energy prediction. The energy dataset was generated to validate the SDDF approach of reducing the need for extensive calculations. With its strict scaffold split, the dataset can be used for training and benchmarking energy models. By combining active learning, distributed computing, and quantum chemistry, SDDF offers a scalable, cost-effective solution for developing accurate molecular models and ultimately accelerating drug discovery.

## 1. Introduction

Machine learning (ML) techniques have reshaped many data-rich fields, such as computer vision and natural language processing, demonstrating that increasing data size can significantly impact model accuracy. This success has sparked interest in applying ML to drug discovery, where data-driven modeling could revolutionize molecular design and property prediction. However, the vast chemical space (“drug-like” chemical space is estimated at the order of 10^60^ molecules [1]) poses a unique challenge: for many critical problems, the available data is limited and often not optimized for machine learning tasks.

Existing molecular datasets, while valuable, present several limitations. Many were not originally designed for training data-hungry ML algorithms but were instead compiled from wet lab experiment reports or created for computational chemistry benchmarks. As a result, their data distribution, coverage of chemical space, and level of theory may not align optimally with the specific requirements of ML methods in drug discovery. For instance, the QM9 dataset [2], widely used for training and benchmarking conformational energy prediction models, provides only one conformation per molecule and lacks scaffold diversity. Furthermore, many of its test set scaffolds (60-99% depending on the split) are present in the training set, indicating data leakage. Similar issues of restricted chemical spaces, potential biases, or limited diversity are seen in other datasets like ANI-1 [3][4][5], NablaDFT [6], and MPCONF196 [7]. All of these datasets provide at best scaffold-based train-test split (without any additional similarity constraints), and also contain mostly smaller molecules relative to what is required for drug discovery projects. Gasteiger et al. 2022 [8] demonstrated the discrepancy in model development decisions, originating from training datasets, and highlighted the importance of dataset diversity and size. Using diverse and large compound libraries of drug-like molecules such as ENAMINE [9] would address most of the aforementioned issues.

The mismatch between existing datasets’ intended use and the emerging needs of modern machine learning techniques extends beyond specific properties like conformational energies. This limitation can hinder the development of robust and generalizable models because the energy landscape of small molecules plays an important role in many other properties, such as binding affinity, solubility, and synthesizability. Therefore, creating high-quality data and methods for energy landscape modeling can be beneficial for various tasks, encompassing a wide range of molecular characteristics crucial for drug design.

Density Functional Theory (DFT) calculations have emerged as a powerful tool for accurately estimating molecular geometries, energies, and various other properties. However, the computational cost associated with these quantum mechanical methods can be prohibitive, especially for large-scale datasets. On the other hand, while machine learning techniques have shown great promise in rapidly predicting molecular properties, their performance is heavily dependent on the quality and diversity of the training data. We provide a more detailed overview of the related work in Supplementary A.

In this paper, we introduce “Smart Distributed Data Factory,” a framework that leverages active learning and volunteer-based distributed computing to construct a comprehensive, high-quality dataset of molecular conformations with their DFT-calculated properties. Volunteer computing has been effectively employed for scientific calculations in many projects before [10][11], including for biochemical problems [12]. With the help of volunteer computing we hope to accelerate these DFT calculations by harnessing the collective processing power of numerous personal computers worldwide. Our approach uses an ensemble of diverse machine learning models to predict conformational energies and strategically selects the most challenging instances for DFT calculations. This targeted selection process ensures that the most valuable data points are prioritized, leading to efficient data acquisition and improved model performance.

Key contributions of our work include:

1. A highly scalable and AI powered distributed computing platform that engages volunteers worldwide to perform QM calculations, accelerating dataset growth.
2. A novel active learning framework tailored to construct a dataset of molecular conformations with DFT-calculated properties, featuring an ensemble of models with different architectures to enhance prediction diversity and sampling efficiency.
3. An ensemble of state-of-the-art machine learning models for accurate prediction of molecule conformational energies.
4. A large and continuously growing public dataset of diverse molecules sampled from the ENAMINE database, that serves as both a training resource and a rigorous benchmark for energy prediction models.

Our methodology offers a scalable and cost-effective solution for building comprehensive datasets tailored to the specific needs of molecular modeling applications. By combining the power of machine learning, distributed computing and quantum chemistry calculations, we aim to enhance the development of accurate and reliable computational models for conformational analysis, ultimately accelerating the discovery and design of novel molecules, particularly for drug development.

The datasets generated in this work are released at zenodo.org/records/14008357. The developed energy prediction models and inference code are freely available at github.com/deeporiginbio/smartdatafactory-experiments.

## 2. Results

We present Smart Distributed Data Factory (SDDF), a distributed computing platform designed for labeled molecule dataset generation using volunteer user machines. It supports a wide range of structure-based molecular property prediction tasks. At its core, SDDF employs an ensemble-based active learning strategy, utilizing models with different architectures to intelligently select the most informative instances from a vast pool of unlabeled data. Having several different architectures is important as the data generation process will select the best architecture for given data size and distribution.

In this section we describe the key elements of the Smart Distributed Data Factory in more detail, including the demonstration of the platform’s use for the creation of a conformational energy dataset. We anticipate that SDDF will be used for the creation of molecule datasets not just for energy predictions, but also other structure-based properties.

### 2.1. Smart Distributed Data Factory volunteer computing platform

The SDDF platform provides a website (https://sddfactory.cloud) where volunteers can sign up and receive molecular conformations for DFT calculations on their personal computers. Each calculation task consists of a single conformation of a molecule and a property specifier indicating a set of properties to calculate. For a targeted property and an average-sized molecule, a single-core machine is expected to calculate the property in about 10 minutes. The result of each task is a dictionary with property names as keys and respective calculated values.

Users can select the projects to which they want to contribute calculations, and they will receive computational tasks only from those projects. Otherwise, the platform assigns tasks from randomly selected projects.

#### Performance benchmarks

To evaluate the performance of SDDF, we conducted several benchmarks. The following aspects were considered:

● Message Broker Reliability: The file-based message broker ensures high reliability and system uptime without the overhead of managing third-party services. Tests indicate that it can handle thousands of messages per second with minimal latency.
● Resource Utilization: Volunteers’ machines use approximately 50% of their computational resources, with each task running on around three threads by default to balance performance and reliability.
● Task Completion Time: For a targeted property and an average-sized molecule (∼25 heavy atoms), a single-core (2.2 GHZ) machine completes a calculation task in about 10 minutes.

By carefully considering these factors, SDDF provides a robust platform for distributed computing, capable of generating high-quality labeled datasets for molecular property prediction.

#### Creating projects for data labeling via DFT

The platform allows to set up different dataset creation projects. For each project, the creator must provide the set of input molecules, and set the parameters for the computational tasks (such as minimum and maximum examples per task, DFT configuration, results verification threshold, etc). The labeled datasets for all projects created via the platform will be publicly available under CC BY 4.0 license.

#### Task definition and volume estimation

We define each computational task as a single molecular conformation for which the user has to calculate the target property using DFT. The volume of each task is estimated based on the number of its conformation’s atoms. The conformations are selected from a pool of unlabeled examples provided by the project’s administrator.

The user side receives the task as an internal gRPC structure, which contains the task type (indicating properties needed to calculate), an identifier and the task content, which is a MOL block representation of desired molecule conformation. After the required calculations are completed, the results are sent to the server side encapsulated in an internal gRPC structure containing the task type, the identifier and the task results represented as a JSON string.

#### Setup and minimal system requirements

The volunteer computing software can be installed on machines with Linux or MacOS systems (support for Windows will be added in the near future). It is set up on the user machines as a Python module. The user needs to specify their credentials and other parameters specified in their accounts in the config file to collect points and climb up in the leaderboard. Then by running the script in the background, the machine starts to periodically fetch tasks from the server, do computations and return results.

To be able to run the computational tasks in a reasonable time, the user machine should be able to allocate 8GB RAM and 10 GB disk space (for storing the temporary files generated during DFT calculations). All computations on the volunteer machines are CPU-only.

### 2.2. Active-learning based data sampling

Smart Distributed Data Factory implements an active learning framework to select molecules for labeling and addition to the dataset. The framework iteratively samples molecules from a large database in random fashion and generates multiple conformations for each molecule using RDKit [13][14] and MD. At each iteration, a fraction of the generated conformations is selected and labeled, after which they are added to the dataset. The selection is performed based on an ensemble of ML models, which are used to determine the most challenging conformations among the generated set of conformations. The target property of the selected most challenging conformations is calculated using DFT. In addition, the selected conformations are used as initial points for MD calculations and the intermediate structures from the calculated trajectories are also labeled via DFT. All newly labeled examples are incorporated into the dataset and used to re-train the ML ensemble. The workflow for the molecule conformational energy dataset creation is illustrated in Figure 1.

**Figure 1.**
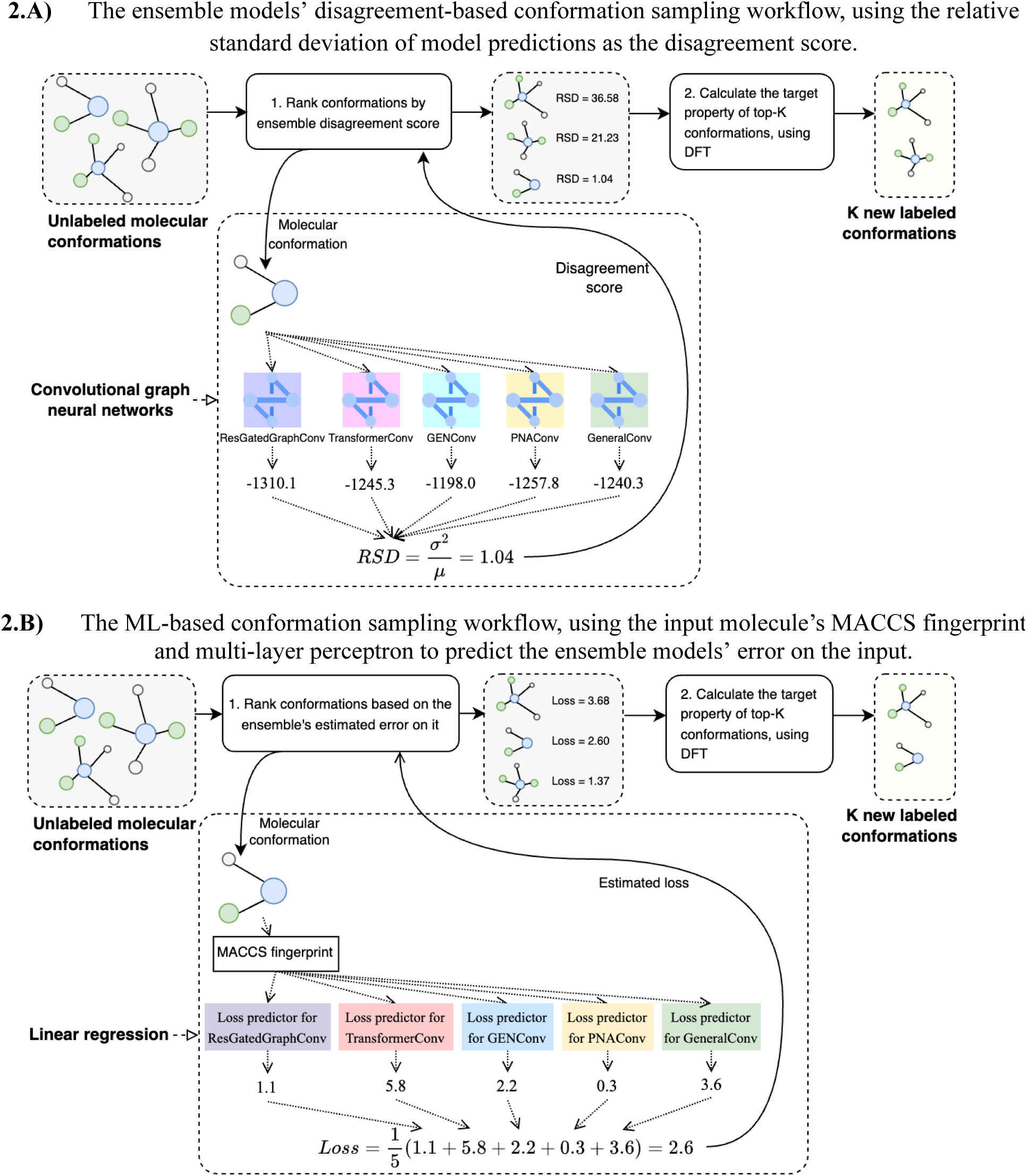
The Smart Distributed Data Factory dataset labeling workflows based on different sampling algorithms: ensemble models’ disagreement-based (2.A) and ML-based (2.B).

In order to train the ML ensemble, our platform labels a small initial dataset of randomly selected conformations, and then its constituent models are re-trained after each iteration of data selection and labeling.

#### Ensemble of predictors

We use an ensemble of ML-based predictors where each predictor is a model trained separately as a regression problem that gets the molecular conformation graph as input and outputs an energy prediction. The nodes of the input graph are the molecule’s atoms and its adjacency matrix is constructed based on the bonds and distances between atoms (we considered an atom pair as adjacent if they have a bond or their distance is below a threshold value).

We performed initial model selection by training and evaluating 33 different graph convolutional neural network (GCNN) and Point Cloud architectures implemented in PyTorch Geometric [15] for the conformational energy prediction task. Based on the evaluation results (Figure 2), we selected the five models with best MAE scores on the validation set: GeneralConv [16], PNAConv [17], GENConv [18], TransformerConv [19] and ResGatedGraphConv [20] models, as implemented in PyTorch Geometric. We further improved the models’ performance by employing Point Pair Features [21] for bonded atoms. The models, their selection and hyperparameter tuning processes are described in more detail in the Methods section and Supplementary B.

**Figure 2.**
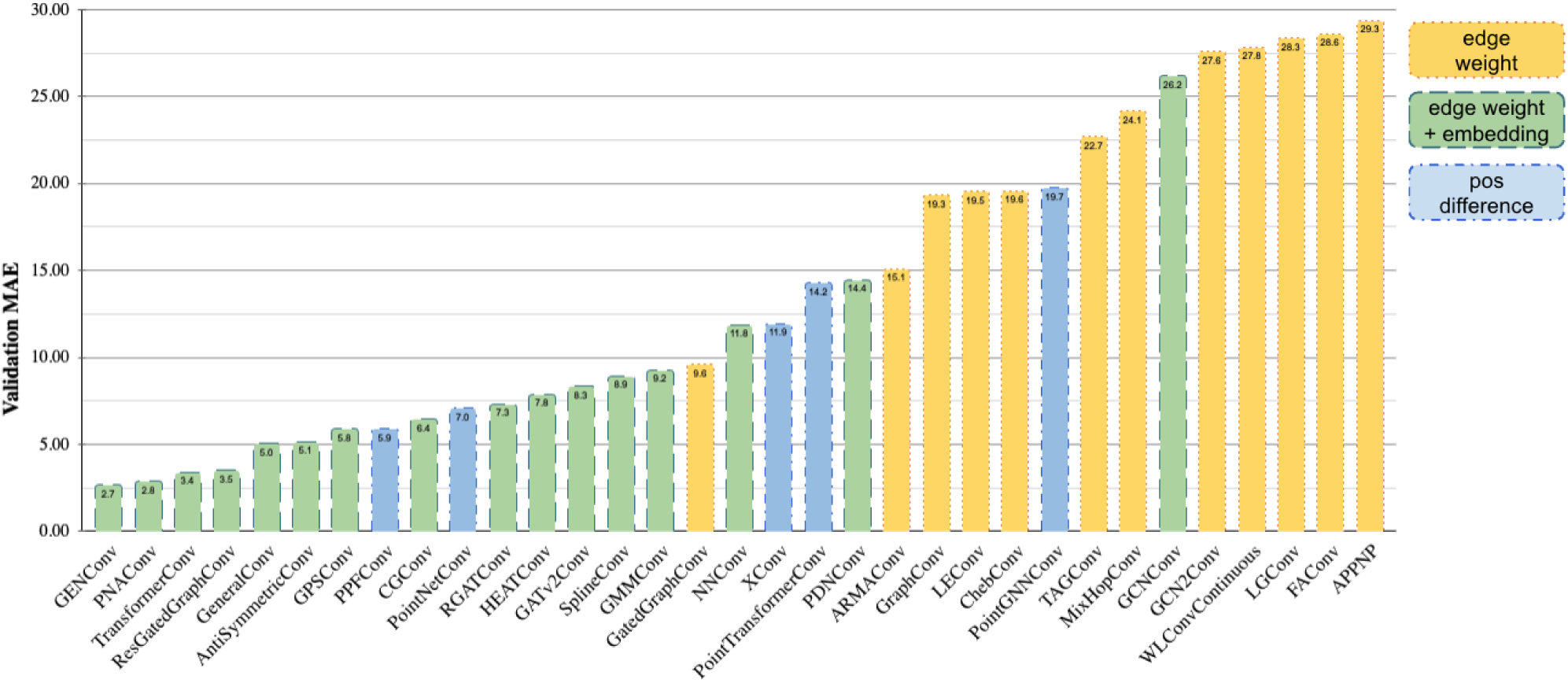
Performance evaluation of GCNN and Point Cloud models on the validation set. (The models are stylized differently based on the method of incorporating the structure’s 3d information: using atom distance-based weights for the edges; using distance-based weights and edge embeddings; using the difference of atom position vectors.)

The decision to use different architectures is based on the assumption that, as the data changes, the effectiveness of the models may also vary. Thus, having a variety of models allows the ensemble to be more robust to distribution shifts in the data. Additionally, we plan to introduce a model selection feature to the platform in the future, where novel architectures could be added to the ensemble if they demonstrate better test scores compared to the existing ones. It is important to note that this selection process will remain dynamic; some models may be dropped if their relative performance declines with more data, while novel architectures will be introduced in their place.

#### Selection of molecular conformations

We investigated different strategies for the selection of molecular conformations for further labeling, with the final approach in our software utilizing machine learning-based predictors. These predictors are linear regression models that estimate the corresponding ensemble models’ error for a given molecule. Each regression model takes the MACCS [22] fingerprint of the molecule as input and is trained on the same dataset as the ensemble. The training process employs a regression approach, predicting the mean absolute error (MAE) between the ensemble model’s prediction and the actual conformational energy. A high predicted error value indicates a challenging and thus valuable instance for data acquisition. The conformations are ranked based on this error (from highest to lowest), and the highest ranked are selected for DFT calculations and subsequent inclusion in the dataset. The predictor’s relatively simple architecture and its independence from 3D structural information enable rapid error estimation across a large molecular dataset, offering an efficient alternative for assessing model uncertainty without relying on variance-based methods.

Additionally, we explored an earlier strategy based on a disagreement score calculated from the predictions of the ML ensemble. For SDDF, this score is based on the relative standard deviation of model predictions, quantifying the uncertainty or conflict among the ensemble’s predictions for a given conformation. While this method has been validated in several previously published works [23][24], the machine learning-based approach ultimately provided a more efficient solution in our final implementation.

To evaluate the effectiveness of the proposed data sampling strategies, we conducted a series of experiments, where we viewed the molecule selection problem in the data stream setup, inspired by data stream active learning techniques [25]. To support our choice of an ensemble with heterogeneous architectures, we investigated four sampling configurations: (i) random sample selection (RAND); (ii) using the relative standard deviation of the predictions of an ensemble of five identically architected GENConv models with different random initializations (x5GENConv); (iii) using the relative standard deviation of the predictions of an ensemble of five different architectural models (SDDF-VAR); (iv) using the model-predicted error of an ensemble of five different architectural models (SDDF-LOSSFN).

Based on the results (Figure 3), we can conclude that using ensemble-based sampling is more effective than random selection. Among the ensemble-based approaches, the most rapid test performance improvement was achieved by the loss prediction-based sampling method. The results also demonstrate the importance of diverse architectures in the ensemble configuration, as SDDF ensemble displayed the most rapid improvement. x5GENConv’s use of varied initializations within the same architecture also allowed for a consistent but slower improvement, highlighting the benefit of using the disagreement of more diverse models. The performance of RANDOM served as a baseline. The experiment results support the hypothesis that ensembles, particularly heterogeneous architectures as in SDDF, can enhance the effectiveness of data selection in active learning, leading to more rapid and potent improvements in model performance compared to homogeneous ensembles or random selection strategies.

**Figure 3.**
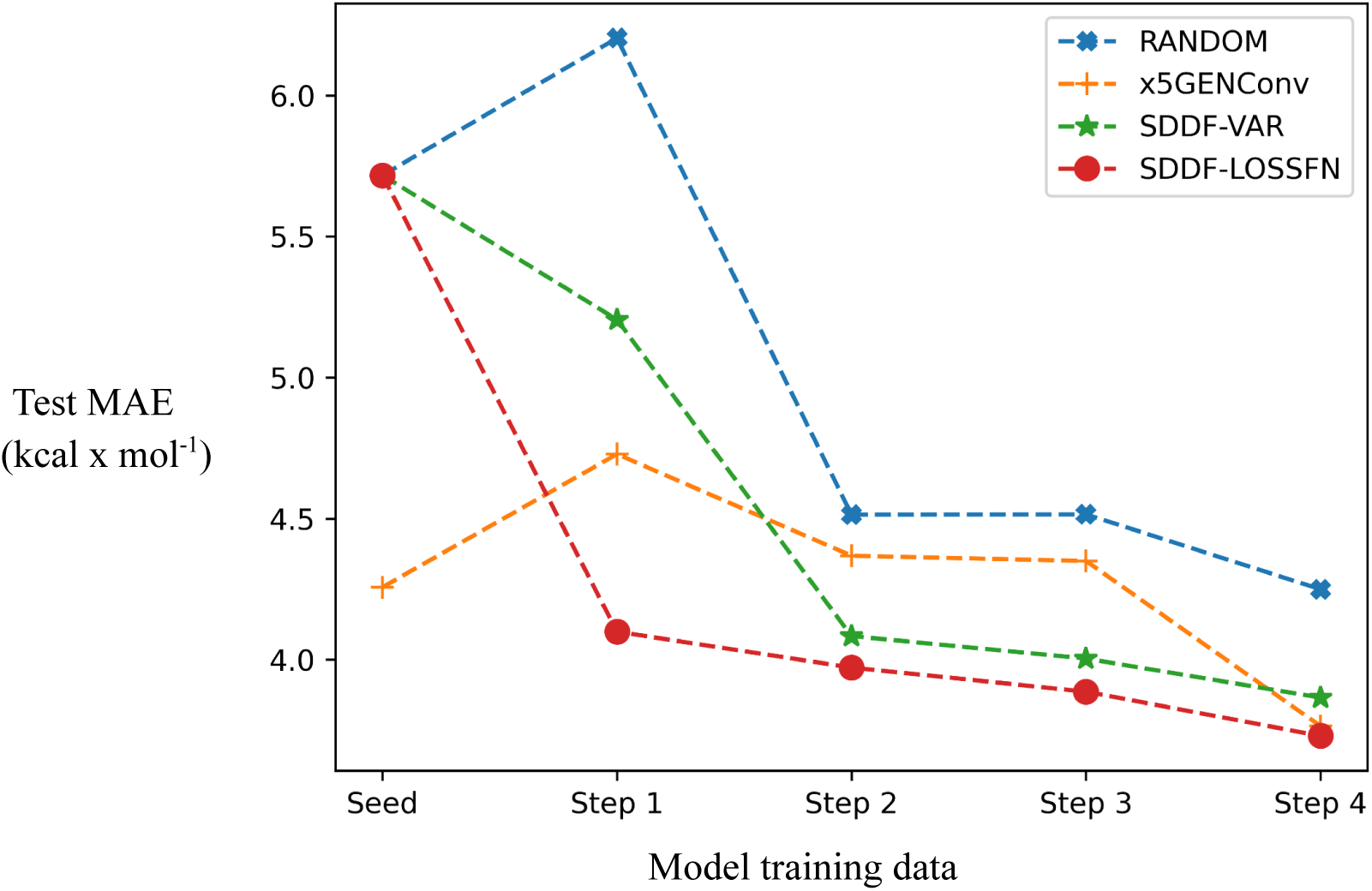
Comparison of data selection strategies: the heterogeneous ensemble-based (SDDF-VAR, SDDF-LOSSFN); the homogenous ensemble-based (5 GENConv with different initializations); random sampling.

### 2.3. Molecular dynamics-based data sampling

In addition to the active learning-based sampling of new data points, we also employ the top-performing models of its ensemble to perform Langevin MD and generate new conformations for labeling. This approach uses a randomly selected small subset of the existing set of conformations, and generates the MD trajectories for each conformation in the subset. For the atoms in the generated conformations we obtain the forces using the gradient of the SDDF ensemble model’s predicted energies. For each atom, we calculate the pairwise cosine similarities of the gradients and select the minimum similarity as a confidence score of the ensemble’s prediction. Then, we compute the average of these atomwise confidence scores for each conformation (Eq. 1), and select the conformations with the lowest average confidence for final labeling via DFT.

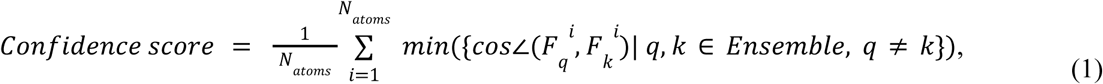

where 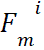 is the gradient of the *m*-th model’s predicted energy with respect to the *i*-th atom’s position.

We evaluated the effectiveness of the MD-based conformation generation and sampling in the task of energy minimization. We randomly selected 1000 conformations from the SDDF test set, and then used SDDF ensembles to generate MD trajectories for these test conformations. MD was performed using 3 different ensembles for the prediction of forces: (i) SDDF ensemble trained on the full SDDF train set, (ii) SDDF+50k-RANDOM ensemble trained on the full SDDF train set and randomly sampled 50k new conformations outside the train the set, (iii) SDDF+50k-MD ensemble trained on the full SDDF train set and additional 50k conformations generated via MD sampling based on 10k random examples from the SDDF train set. Thus, we obtained 3 new sets of conformations and labeled them using DFT. We calculated the difference in the energies for the conformations in the 3 generated conformation sets and the original set to estimate the impact of additional data on the ensemble’s ability to guide the molecules to a lower-energy conformation.

We calculated the portion of conformations where the energy decreased after MD, and the average difference between resulting conformations’ energies and the original energies (Table 1). The results show that the addition of training examples generated via MD-based sampling has a much more positive impact on the ensemble’s ability to do MD than the addition of randomly sampled train examples of the same number. The generally low success rate in minimizing energy may be partly due to the fact that the initial conformations generated by RDKit had already been optimized with the MMFF94 force field before these MD steps.

**Table 1.** Comparison of SDDF ensembles in the conformational energy optimization task. Success rate is measured as the portion of original conformations for which the energy was minimized during the MD.

|  | SDDF | SDDF+50k-RANDOM | SDDF+50k-MD |
| --- | --- | --- | --- |
| Success rate | 8.42% | 9.25% | **26.32%** |
| Minimized energy average | 0.0084 | 0.0091 | **0.0016** |

### 2.4. Conformational energy datasets and machine learning ensemble

#### Conformational energy dataset and benchmark

Smart Distributed Data Factory (SDDF) relies on a relatively small starting dataset of labeled examples, which are used to train its ensemble’s initial models. For the conformational energy prediction task, we create and publish a dataset of 2.17M molecular conformations with DFT-calculated energy labels. The molecular conformations were generated from ENAMINE molecules, using Python’s RDKit toolkit with different random seeds. DFT calculations were performed through Python’s Psi4 toolkit, using the same theory level as ANI datasets: ωB97x [26] density functional and the 6–31 G(d) basis set [27]. We selected the REAL database of ENAMINE as the source of our examples, because it is a pivotal and extremely relevant collection of synthetically possible molecules used for drug discovery and development projects.

Our full labeled dataset contains 2,170,553 conformations, including 535,338 that were generated using RDKit, another 1,151,936 that were generated using RDKit and optimized with the MMFF94 force field, and 483279 other conformations generated using MD on RDKit conformations.

In addition, we release a subset of the dataset for training and benchmarking energy prediction machine learning models (the statistics of the split are available in Supplementary C, Table 14). We applied a strict scaffold split to our training, validation and test sets, so that the maximum Tanimoto similarity between any test and train scaffolds did not exceed 0.7. In this work, we determine the scaffolds using the RDKit implementation of the Bemis-Murcko framework [28].

Apart from using ENAMINE as the source of the molecules, the decision to use a new dataset for the energy prediction task was motivated by other important factors as well: the lack of either conformational or scaffold diversity (Supplementary C, Table 13), the relatively small size of the molecules in the existing datasets (Supplementary C, Figure 11), or the presence of train test leakage. For example, the QM9 dataset, which is often used as a benchmark, does not provide more than a single conformation for each molecule, and at the same time lacks scaffold diversity (the overall ratio of unique scaffolds to total examples is less than 2%). In addition, more than 60% of the test set scaffolds (Cormorant dataset split [29]) are also present in the train set, indicating leakage from that set.

#### Comparison with ANI-2x

Using the architectures selected for the active learning framework, we trained conformational energy prediction models using our training and validation sets. Alongside the datasets, we also release these models.

In order to validate the effectiveness of the selected model architectures in the energy prediction task, we compared their performance with the ANI-2x ensemble [30]. Table 2 provides the evaluation results of the models on our test set. We used the TorchANI toolkit (version 2.2.4) [31] to perform the inference for ANI-2x. Our ensemble and the majority of its individual models outperformed the ANI-2x ensemble in RMSE and MAE metrics.

**Table 2.** Performance of ANI-2x and SDDF ensemble models on our test subset without Bromine atoms (scores in the brackets indicate the full test set MAE in kcal*mol^-1^, including examples with Bromine atoms).

| Model |  | Test RMSE* |  | Test MAE* |  |
| --- | --- | --- | --- | --- | --- |
| SDDF | PNAConv | 3.38 | (3.35) | 2.06 | (2.06) |
|  | ResGatedGraphConv | 3.42 | (3.39) | 2.24 | (2.25) |
|  | GENConv | 3.60 | (3.58) | 2.23 | (2.25) |
|  | GeneralConv | 5.97 | (6.02) | 4.08 | (4.10) |
|  | TransformerConv | 7.34 | (7.31) | 5.11 | (5.12) |
|  | Ensemble{PNAConv, ResGatedGraphConv, GENConv} | **2.92** | **(2.89)** | **1.82** | **(1.83)** |
|  | Ensemble{all} | 3.54 | (3.52) | 2.30 | (2.31) |
| ANI-2x* |  | 5.07 | — | 2.84 | — |

\*Does not support Bromine

We additionally analyzed the dependence of the models’ performance on the size of test molecules (Figure 4). The ANI-2x ensemble shows noticeably higher MAE on molecules outside its train data distribution (> 25 atoms), while the MAE of the SDDF ensemble remained relatively more stable across different molecule sizes (within 5 MAE, with the exception of molecules with 20-25 and 55-60 atom counts for SDDF ensembles).

**Figure 4.**
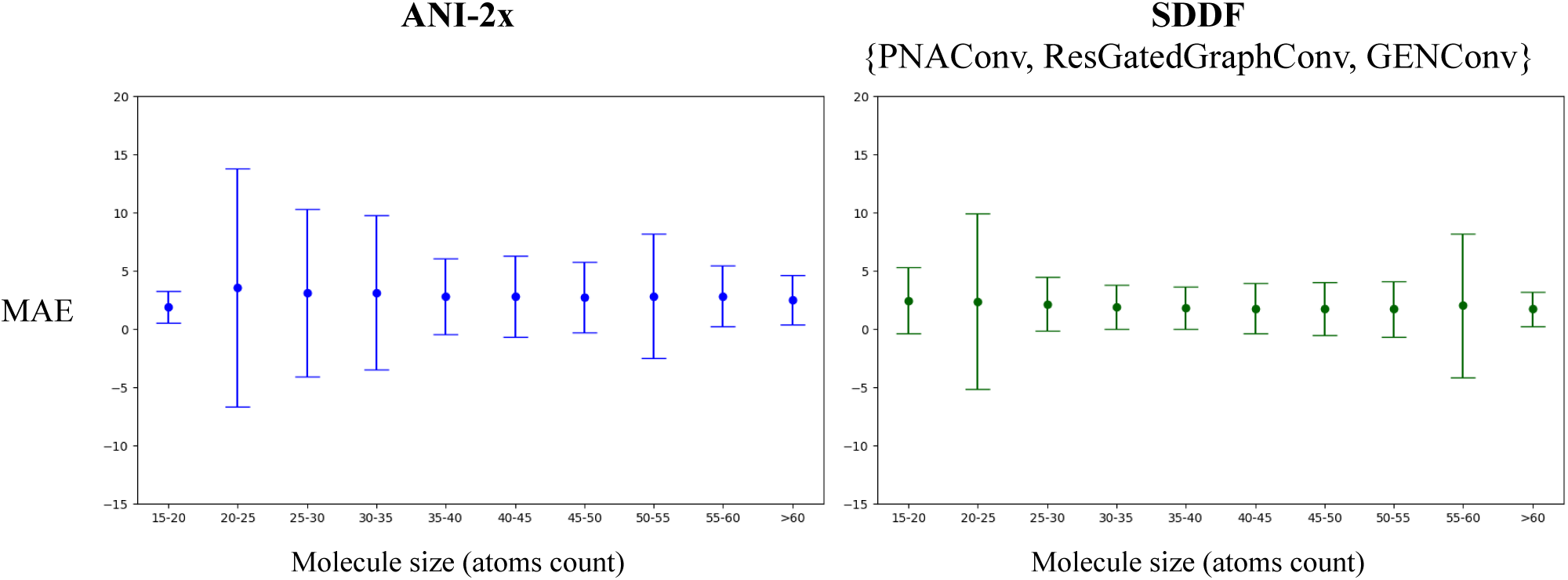
Comparison of models’ test set prediction errors based on molecule size.

## 3. Discussion

In this paper, we introduced Smart Distributed Data Factory, a framework for generating labeled datasets in computational chemistry. Our ensemble-based active learning strategy, which estimates the error of machine learning models on individual instances, efficiently identifies molecular conformations where high prediction errors are expected, thus prioritizing those instances for DFT calculations.

The Smart Distributed Data Factory framework integrates two essential components: an ensemble of machine learning models for data selection and a volunteer computing platform for performing DFT calculations. The ensemble consists of five architecturally distinct graph neural networks, which improves its ability to capture a diverse range of molecular complexities and identify instances likely to result in higher errors. This approach ensures that the most challenging and uncertain data points are selected for labeling. Simultaneously, the volunteer computing platform distributes computational tasks to volunteers globally, enabling scalable and distributed data labeling, which significantly enhances the framework’s capacity.

By applying these methods, we created a dataset of molecular conformations along with their DFT-calculated energies. This dataset serves as both a valuable training resource and a benchmark for evaluating energy prediction models. By focusing on instances where models estimate higher prediction errors, the active learning strategy ensures that the dataset is rich in challenging examples, enhancing the overall quality of the training data.

However, several challenges remain. The scalability of the volunteer computing platform is inherently dependent on participant availability and resource distribution, which may result in uneven computational capacity. Sustaining volunteer engagement and optimizing task distribution algorithms are areas for future improvement. In addition, the platform currently provides computational projects for only conformational energy and atomic charges calculations. In the future, we plan to add projects for the calculation of other properties. Furthermore, while the ensemble of graph neural networks has proven effective, refining active learning strategies and incorporating additional architectures could also improve the framework’s performance. This includes transitioning from fingerprint-based linear regression to more complex methods for predicting the ensemble’s error, such as using 3D structure-sensitive models (for example, GCNNs).

In conclusion, Smart Distributed Data Factory demonstrates the potential of combining active learning with distributed computing for efficient dataset creation in computational chemistry. By focusing on instances with high error estimates, this framework contributes to the development of more accurate and reliable models for molecular property prediction.

## 4. Methods

### 4.1. Smart Distributed Data Factory system architecture

The main components of the Smart Distributed Data Factory system implementation (illustrated in Figure 5) are the following:

● Central Node:
  ○ Task Queue: Receives tasks from the Task Publisher and manages the queue of tasks to be processed.
  ○ Database: Stores data necessary for task processing and system management.
  ○ SDDF Server: Responsible for forming computational tasks and their distribution. Communicates with the Central Node for data exchange via gRPC.
  ○ Web Server: Hosts the website and handles interactions with external clients.
● Distribution Node:
  ○ SDDF Tunnel: Facilitates the transfer of computational tasks and results between the Distribution Node and the Central Node using gRPC.
  ○ SDDF Client: Volunteer nodes connected to the Distribution SDDF tunnel for getting molecular structures to perform computational tasks.
● Scheduled Services:
  ○ Molecular Conformation Generator: Generates 3D conformations from molecular SMILES using either RDKit implementation of the ETKDGv3 algorithm [13], or Openbabel [32], optionally optimizing generated conformations using MMFF94 force field.
  ○ ML-Based Force Field Conformation Generator: Generates new conformations by running molecular dynamics starting from some existing conformation. Instead of traditional force-fields it uses negated gradients of our energy predicting models with respect to coordinates as atomistic forces.
  ○ AI Enhanced Task Selector: Selects from the generated set of conformations the ones which are challenging for our existing models and pushes them to the task queue. The web server is a FastAPI-based leaderboard system designed to track and display volunteer contributions. It uses a MongoDB database for storing and retrieving user data and contributions.

**Figure 5.**
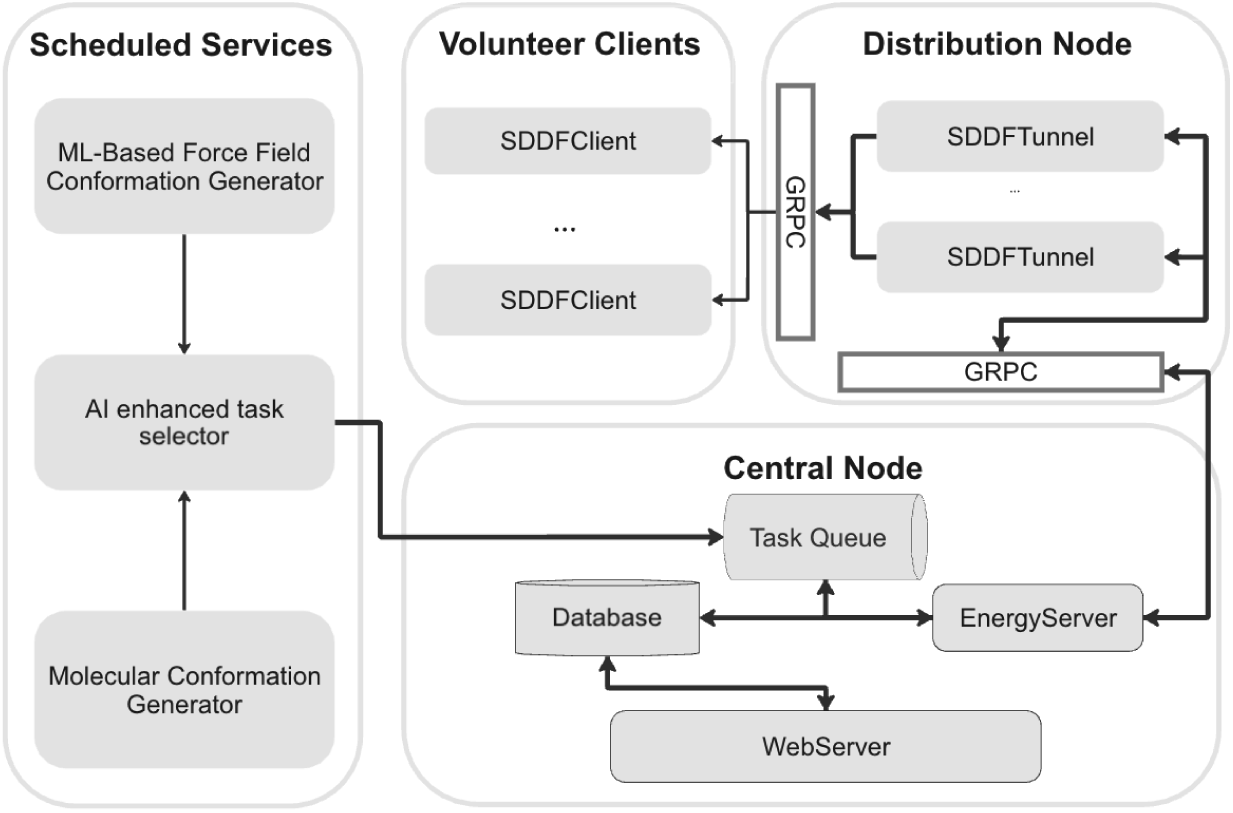
The architecture of the distributed computing system.

To handle task queuing and distribution, the platform uses a file-based message broker. This approach leverages external volumes for storage, ensuring reliability and persistence of messages. A primary benefit of this design is avoiding the overhead of managing third-party services, thereby contributing to system uptime and simplifying maintenance. Comparative analyses with third-party message brokers like RabbitMQ and Apache Kafka demonstrate that while these services offer robust features, they also come with additional management overhead. The file-based message broker provides a lightweight, reliable alternative suitable for environments prioritizing uptime and simplicity. gRPC is utilized for communication between different nodes in the system.

The SDDF Client computations are CPU-only, and can be run on architectures supported by the Psi4 library for quantum chemistry calculations.

### 4.2. Smart Distributed Data Factory ensemble model selection and simulation strategies

#### Initial labeled data

Smart Distributed Data Factory (SDDF) relies on a relatively small starting dataset of labeled examples, which are used to train its ensemble’s initial models. For the conformational energy prediction task, we obtained our initial dataset by calculating energy for around 800k molecular conformations using DFT. The molecular conformations were generated from ENAMINE molecules, using Python’s RDKit toolkit with different random seeds. DFT calculations were performed through Python’s Psi4 toolkit, using ωB97x density functional and the 6–31 G(d) basis set.

#### Train, validation, and test split

From the initial labeled data we obtained training, validation, and test sets using the approach below:

1. We randomly selected 80% of the molecules as the training set.
2. From the other 20% we selected as the validation set the molecules that had 70% or higher scaffold similarity to any training molecule.
3. The remaining molecules made the test set.

This approach generates splits that meet the following criteria:

1. No matching molecules between the sets.
2. The maximum scaffold similarity between any pair of molecules in the training and test sets should be lower than 70%.

The generated test set contains nearly 25k conformations with over 6k unique scaffolds that do not overlap with the train set examples.

#### Data stream simulation setup

For SDDF evaluation we simulate a setup of active learning on unlabeled data stream. Based on the train set, we initialize 4 sets of examples:

● **Pool:** the set of examples, from which data stream is simulated.
● **Buffer:** the set of examples which are ranked to select labeling candidates.
● **Seed**: the set of examples for training the ensemble’s initial models.
● **SDDF**: the continuously growing set of examples selected via Smart Distributed Data Factory algorithm that have corresponding DFT calculations.

We initialize Seed as 100k randomly picked conformations from the train set. From the rest of the train set, Buffer is initialized with 60k random examples. Then, Pool is populated with the train set examples that were not included in either Seed or Buffer. SDDF is empty at the start of simulation.

The evaluation consists of several data selection rounds. During each round we re-train the ensemble models on the combined Seed and SDDF sets, then use the updated ensemble to re-rank Buffer examples based on a disagreement metric, and move the top-ranked 30k examples from Buffer to SDDF. Finally, we re-populate Buffer with 30k random examples from Pool (these examples are removed from Pool). The process is illustrated in Figure 6. To reduce the impact of instability caused by random initialization in the GCNN models’ performance and to obtain a reliable estimate of the data selection strategies’ effectiveness, we ran the simulations using different random seeds and averaged the results.

**Figure 6.**
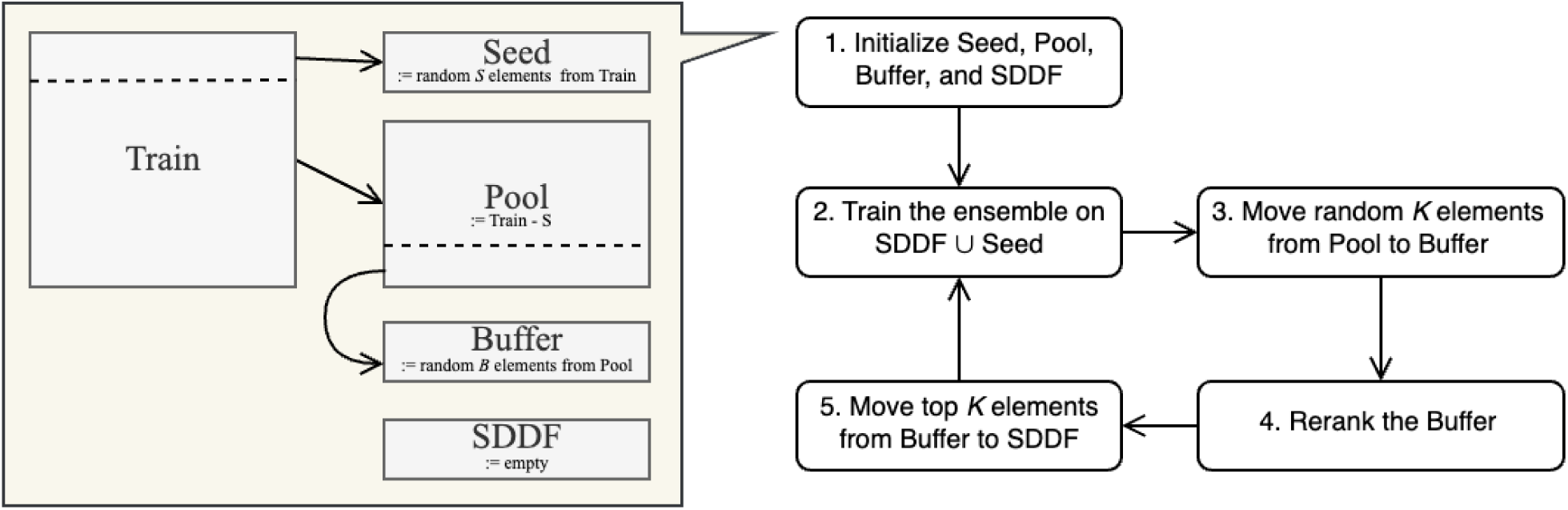
Smart Distributed Data Factory simulation setup.

#### Ensemble model selection

In order to select appropriate neural network architectures for use in the ensemble of Smart Distributed Data Factory, we studied the PyTorch Geometric implementations of more than 40 GCNN and Point Cloud model architectures from prominent ML conference papers. Specifically, we assessed the ability of methods to incorporate structure information based on the the presence of at least one of the following features:

I. Usage of edge weights during message passing. This would allow us to incorporate the molecular conformation information by using distance-dependent weighting for messages.
II. Usage of edge embeddings during message passing. We can encode atom pair distance and other 3d information into edge embeddings.
III. Usage of atom pair position difference. Position difference is an input feature in some message passing networks, and some attention-based models use it during attention calculations. Thus, the predictions of these models are also conformation-dependent.

We picked 33 appropriate candidates and separately trained each candidate architecture on our train set and evaluated on the validation dataset, using the Adam optimizer and Mean Absolute Error as the optimization loss and validation metric. The details of input features and training hyperparameters are provided in the Supplementary B.

### 4.3. Molecular dynamics settings

When evaluating the effectiveness of adding MD generated conformations to our datasets, we created train and test examples. For generating training data, the MD simulation for each starting conformation was run for up to 1000 steps (1 picosecond), and every 10th step was saved to a trajectory file. For each starting conformation at most 6 new conformations were selected with lowest values of the ensemble’s confidence score for the predicted forces. As an early stopping criterion, we set a constraint on the system temperature, approximately calculated from the system’s kinetic energy and the number of degrees of freedom.

The MD simulation for the test set of this experiment was run for at most 100 steps (0.1 picoseconds) and each step of the simulation was saved to a trajectory file. The early stopping criterion was applied here too, and the upper bound for temperature during simulation was chosen as 600K. This was done in order to prevent the system from breaking upon the simulation start, as velocities are initialized randomly.

## Data Availability

The energy datasets generated during the current study are available in the Zenodo repository “SDDF Energy Dataset”, https://doi.org/10.5281/zenodo.14008357.

Other datasets analyzed during the current study:

● **ANI-1** is available in the Figshare repository “ANI-1: A data set of 20M off-equilibrium DFT calculations for organic molecules”, doi.org/10.6084/m9.figshare.c.3846712.v1.
● **ANI-2x** is available in the Zenodo repository “ANI-2x Release”, doi.org/10.5281/zenodo.10108942.
● **NablaDFT** analysis was performed using the summary file provided in the GitHub repository’s README.md file (github.com/AIRI-Institute/nablaDFT/blob/1.0/README.md). Persistent web link to the summary file: a002dlils-kadurin-nabladft.obs.ru-moscow-1.hc.sbercloud.ru/data/nablaDFT/summary.csv.
● **NablaDFT 2.0** analysis was performed using the summary file provided in the GitHub repository’s README.md file (github.com/AIRI-Institute/nablaDFT/blob/main/README.md). Persistent web link to the summary file: a002dlils-kadurin-nabladft.obs.ru-moscow-1.hc.sbercloud.ru/data/nablaDFTv2/summary.csv.gz.
● **GEOM** is available in the Harvard Dataverse repository “GEOM”, doi.org/10.7910/DVN/JNGTDF.
● **QM9** is available in the Figshare repository “Quantum chemistry structures and properties of 134 kilo molecules”, doi.org/10.6084/m9.figshare.c.978904.v5.
● **MD17** and **MD22** datasets are available in the sGDML.org webpage, www.sgdml.org/#datasets.

## Code availability

The source code of the machine learning model training and evaluation infrastructure is available at <u>github.com/deeporiginbio/smartdatafactory-experiments</u>. The source code for the volunteer computing platform is available to researchers upon reasonable request. Please contact the corresponding author for access.

## Acknowledgements

We would like to thank all the volunteers who contributed their computing resources to this project. We are also grateful to our colleagues at Deep Origin for their valuable feedback and assistance throughout the course of this work.

## Author contributions statement

● Vahagn Altunyan designed and implemented the volunteer computing platform and contributed to the active learning framework. He prepared the datasets, generated molecular conformations, and labeled them via DFT. He also contributed to the experimental design and the manuscript writing.
● Garik Petrosyan contributed to the design of the volunteer computing platform and the active learning framework. He also contributed to the experimental design, supervised the overall project, and revised and reviewed the manuscript.
● Tsolak Ghukasyan designed the experiments and participated in the development of the machine learning models and the infrastructure for testing the active learning framework. He analyzed and interpreted data from benchmarks and model testing, drafted the manuscript, and prepared figures and supplementary materials.
● Tigran Aghajanyan and Aram Bughdaryan were the main developers of the infrastructure for testing the active learning framework and model training. They performed the analysis and selection of machine learning models, handled the data splits, and prepared figures and supplementary materials.
● Khachik Smbatyan contributed to the design and development of the platform.
● Garegin A. Papoian provided scientific guidance for the project and contributed to manuscript review and revision.

## Additional information

The authors of this paper are employees of Deep Origin Inc. and hold shares of the company. The authors have no other relevant affiliation or financial involvement with any organization/entity that has a financial interest in or financial conflict with the subject matter and materials discussed in this paper.

## Supplementary A. Related work

The use of machine learning (ML) techniques for predicting molecular properties, including conformational energies, has been an active area of research in recent years. One of the key challenges in developing accurate ML models for molecular modeling applications is the availability of high-quality training data.

### Chemical space sampling for dataset creation

Traditional approaches to building datasets often rely on brute-force sampling or random selection of molecules, which can be computationally expensive and may fail to capture the most informative instances. To address this issue, several researchers have explored various active learning strategies for intelligent data acquisition.

● Artrith and Behler et al. (2012) [33] performed a neural network-based MD simulation and then used another network to predict the potential energies of the generated trajectory’s conformations. They then used DFT to re-calculate the potential energies for the conformations where the predictions of the networks differed from each other, and used these conformations with the newly calculated energies as training examples in the future.
● Zhang et al. (2019) [24] presented the deep potential generator DP-GEN, based on an ensemble of Deep Potential models. They used the maximum standard deviation of predicted atomic forces as the criteria for selecting examples for DFT labeling.
● Smith et al. (2018) [23] proposed using an active learning framework based on an ensemble of ANI models for the creation of a labeled conformational energy dataset, emphasizing that “Less is more” when it comes to dataset size versus quality. In their setup, they iteratively select and calculate potential energies for samples from a given fixed set of conformations until the performance of the ensemble reaches the pre-defined threshold.
● Jung et al. (2024) [34] also used an ensemble of ANI models for active learning-based data sampling, additionally using similarity checks for select unique configurations and augmenting data by adding random noise displacement at each active learning iteration.

Our work builds upon these previous efforts but has several key differences:

1. As the source for molecules for the datasets we use ENAMINE, which is an essential database for drug discovery projects.
2. We performed a comprehensive evaluation of graph neural-network based models and selected an ensemble of distinct models with different architectures. The heterogeneous ensemble approach helps to mitigate the biases of individual models and improves the overall performance of the data sampling process.
3. We performed evaluation of AL sampling methods and selected best performing selection method based on ML models
4. We employ molecular dynamics to further enrich the set of molecular conformations.
5. We introduce a volunteer computing platform, distributing selected sets of molecule conformations to volunteers to perform the calculations on their machines.

### Volunteer computing

In creating our system, which uses volunteer computing to analyze molecular properties via QM, we reviewed various distributed computing platforms. Although each provided valuable insights, none

completely suited our project’s unique needs, especially the need for AI-enhanced datapoint sampling, prompting us to develop a new platform.

● Globus Toolkit (Foster and Kesselman, 1997) [35] and Grid Services for Distributed Systems Integration (Foster et al., 2002) [36] have pioneered in providing secure, high-performance distributed computing infrastructures and service-oriented architectures, respectively. These frameworks excel in academic and institutional settings but lack the flexibility and accessibility needed for engaging a wide, public audience in computationally intensive DFT calculations.
● SETI@home (Anderson et al., 2002) [37], Cosm [38], JXTA [39][40], and XtremWeb (Fedak et al., 2001) [41] demonstrated the potential of public-resource and peer-to-peer computing for tackling large-scale scientific problems through volunteer participation. However, these platforms are either too specialized (SETI@home) or not optimized (Cosm, JXTA, XtremWeb) for the specific, high-precision computational tasks required in chemistry, such as DFT analysis.
● Entropia (Chien et al., 2003) [42] offered utilizing idle computing resources within more controlled environments, such as corporate networks or through commercial platforms. While they showcased the viability of distributed computing for scientific and commercial applications, their models do not align with our vision of an open, collaborative, and freely accessible platform designed for the global scientific community and public volunteers.
● BOINC (Anderson, 2004) [10][11] stands out for its framework supporting volunteer computing across various scientific disciplines, closely aligning with our objective of widespread public engagement. There have been several usages of BOINC for biomedical research purposes, such as GPUGRID.net [43] for all-atom biomolecular simulations and Folding@home [12] dedicated to understanding protein folding. However, its general-purpose nature requires considerable adaptation to meet the nuanced demands of machine learning model training and inference, molecular property computations using DFT, including task specificity, computational efficiency, and the management of complex scientific data.

Consequently, while acknowledging the foundational work of these predecessors, the unique challenges of our project—namely, the need for a highly flexible, accessible, and scientifically rigorous AI enhanced computing platform—necessitate the development of a solution tailored to the dynamic and precise requirements of molecular simulations and DFT calculations.

### Conformational energy datasets

Several publicly available datasets contain energy annotations for molecular conformations, but each presents its own limitations or drawbacks:

● ANI-1 [4] dataset is based on a subset of the GDB-11 database. It contains DFT-calculated energies for approximately 20 million conformations of small organic molecules. However, it is limited to molecules with only H, C, N, and O atom types, with a maximum of 8 heavy atoms (C, N, O). This restricted chemical space may not be representative of the diverse molecular structures encountered in drug discovery or material science applications.
● ANI-2x [30], a more extensive and diverse follow-up dataset to ANI-1, this dataset provides DFT properties at 5 different levels of theory for small organic molecules containing H, C, N, O, S, F, and Cl atom types. While large in the number of molecules and inclusive of more atom types, this dataset mainly contains relatively small molecules.
● NablaDFT [6]: Introduced by Khrabrov et al. 2022, this large-scale dataset comprises 6 million conformations for 1 million molecular structures with C, N, S, O, F, Cl, Br, and H atom types. Although it provides broader coverage of chemical space compared to ANI-1, the dataset is derived from a subset of the MOSES dataset, which was initially designed for molecular generation tasks and may not be optimized for conformational analysis or energy prediction. The conformations in NablaDFT were generated using RDKit [13], without employing molecular dynamics (MD) or sophisticated conformation sampling algorithms. In their recent follow-up work, NablaDFT 2.0 [44], they doubled the number of molecules and conformers in the dataset, and also released approximately 3 million conformations obtained from the relaxation trajectories for around 60,000 examples.
● GEOM [45] dataset contains 37M energy-annotated molecular conformations for 133,000 different molecules from QM9 and 317,000 molecules from experimental data related to biophysics, physiology, and physical chemistry. The average number of heavy atoms in the molecules is below 20 (although if we exclude QM9 molecules, the average rises to over 25, which is a bit higher than NablaDFT), and the level of theory used for DFT calculations is considered by some [44] to be relatively less accurate than the theory level of ANI and NablaDFT datasets.
● MPCONF196 [7]: This benchmark dataset focuses on accurate conformational energies of smaller peptides and medium-sized macrocycles. While valuable for its specific domain, the dataset may not be representative of the broader chemical space relevant to drug discovery or material science applications.
● QM9 [2]: Created by researchers at the University of Warwick, the QM9 dataset consists of geometric, energetic, and electronic properties for a subset of 133,885 stable and synthetically accessible organic molecules, comprising up to 9 heavy atoms (C, N, O, F). While widely used for benchmarking, the dataset’s coverage is limited to a specific range of molecular sizes and atom types. Additionally, as noted in the introduction, QM9 provides at most one conformation per molecule, and its overall ratio of unique scaffolds to total examples is less than 2%. Moreover, more than 99% of its test set scaffolds are also present in the train set, indicating leakage.
● QMugs [46]: This dataset, developed by researchers at the University of Cambridge, contains quantum mechanical properties, including energies, for a diverse set of molecular structures. However, the specific details regarding its composition, diversity, and potential biases for conformational analysis tasks are not widely documented or evaluated in the literature.
● Transition1x [47]: This dataset contains 9.6 million DFT calculations of forces and energies of molecular configurations on and around reaction pathways.
● MD17 [48]: The dataset is a collection of over 3.5 million MD trajectories for 8 small organic molecules. MD22 [49], regarded as the next generation of MD17, contains over 200,000 MD-simulated conformations for 7 systems with 42 to 370 atoms.

In our datasets we include molecules from the ENAMINE database, with C, N, S, O, F, Cl, Br, and H atom types. We generate conformations via RDKit, but additionally obtain new conformations using MD. We also provide a train and test benchmark with strict scaffold split.

### Conformational energy prediction with machine learning

In recent years, several neural network architectures have emerged which allow the creation of conformation aware models. It is fundamental for all methods to be invariant to translations and rotations of the input molecule’s coordinates. ANI-1 model and its extensions [5][30][31] use multi-layer perceptrons and single-atom atomic environment vectors as input for neural networks-based energy and force prediction in molecular systems. Graph convolutional neural networks (GCNN) have been a particularly popular architecture for molecular property prediction tasks [50] and still demonstrate competitive results on several benchmarks [51][52][53][54]. Transformers [55] have also shown great capacity to encode 3D structural information via adapted attention mechanisms [56], and have been adopted in several molecular property prediction models [57]. More recently, SE(3)-equivariant Transformers have gained popularity [58][59][60][61], showing state-of-the-art results on some benchmarks.

## Supplementary B. Smart Distributed Data Factory platform

This section describes the capabilities and implementation of our volunteer computing platform in more detail, including the ensemble of machine learning learning for active learning-based data sampling, their architecture and training hyperparameters.

### Platform implementation details and features

gRPC is utilized for communication between different nodes in the system. gRPC offers several advantages, including:

● High Performance: gRPC uses HTTP/2 for transport, providing efficient binary serialization and reducing latency. Benchmarks [42] indicate that gRPC can achieve significantly better performance in terms of latency and throughput compared to traditional REST APIs. For example, gRPC handled requests almost seven times faster than REST in a benchmark test, with gRPC achieving 141 requests per second compared to REST’s 22.9 requests per second.
● Scalability: gRPC supports multiple concurrent connections and efficient load balancing, crucial for the system’s scalability. Google’s internal use of gRPC demonstrates its capability to handle large-scale, distributed systems efficiently.
● Reliability: gRPC’s strong typing and contract-first approach ensure reliable and well-defined communications between nodes, reducing the risk of errors. It includes built-in support for automatic retries, backoff strategies, and deadlines/timeouts, enhancing the reliability of communications.

In order to verify the calculation results and also to prevent the malicious use of the platform by a single user or a group of users, the submissions by all users are continuously screened. Upon registration, each user first receives several tasks, for which the computation results are already known. The submitted results are checked against the known results, and if they do not match for 80% of the cases, the user is not sent further tasks. After the initial screening procedure, the submissions of the user are continued to be verified at a rate of 1 per 100 tasks on average.

The platform’s website also provides some community features such as a public leaderboard of top-ranked contributors. The ranking is based on the estimated amount of total calculations by each user. Visitors can also apply different filters on the leaderboard and, for example, view the leaderboard for each project.

### Ensemble of machine learning models for active learning

In this section, we provide a detailed explanation of our ensemble model selection process and the performance of the machine learning models used for active learning and energy prediction tasks. The ensemble approach combines various graph convolutional neural networks (GCNNs) models, each with different ways of aggregating messages and applying attention. We explain the structure and function of the selected models GeneralConv, PNAConv, GENConv, TransformerConv, and ResGatedGraphConv. To improve accuracy, we used node and edge representations, including Point Pair Features (PPF), which helped boost the performance of molecular energy predictions. We also compare our results with leading models like ANI-2x and GemNet, focusing on how they perform with molecules containing bromine atoms. The results show that even without using ensembles, many of our individual models outperform ANI-2x. The following sections contain analysis on how our models performed during training, validation, and ranking tasks, along with a look at the distribution of molecules selected in an active learning simulation.

#### Ensemble model selection

We trained and evaluated 33 different neural network models implemented in PyTorch Geometric [15] for predicting conformational energy. From these evaluations (Table 3), we chose the top five models with the best MAE scores on the validation set: GeneralConv [16], PNAConv [17], GENConv [18], TransformerConv [19], and ResGatedGraphConv [20], all implemented in PyTorch Geometric. We also demonstrated that incorporating Point Pair Features [21] for bonded atoms enhances the performance of the models.

**Table 3.**
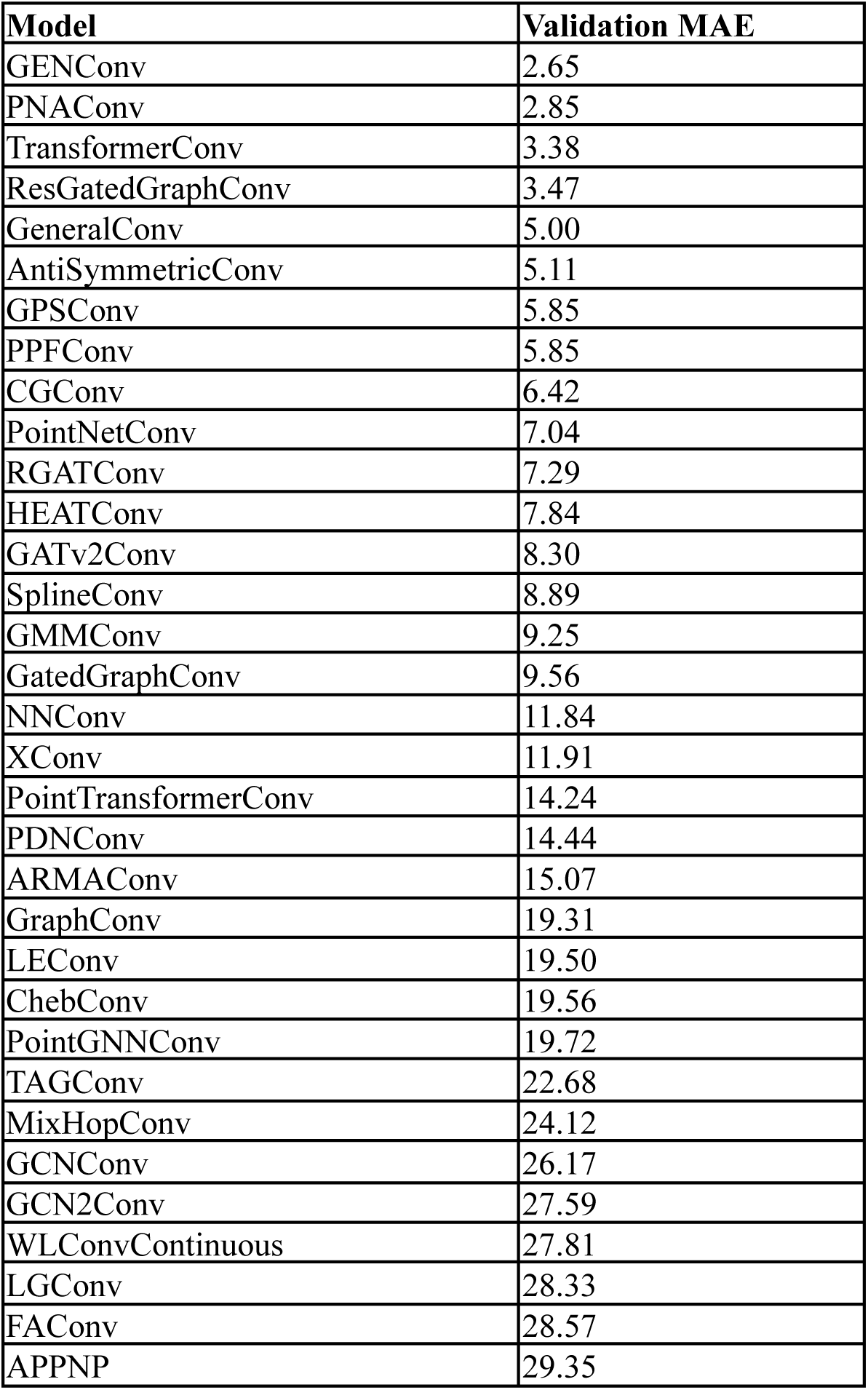
Performance evaluation of GCNN and Point Cloud models on the validation set.

GeneralConv is the implementation of the general GCNN layer from the “Design Space for Graph Neural Networks” [16] paper. We used the mean aggregation scheme for the messages, with only 1 message calculated per atom pair, without linear function in skip connection and without adding attention to message computation. We included the edge embedding in the computation, weighting each embedding based on the distance between the atoms.

PNAConv architecture from the “Principal Neighbourhood Aggregation for Graph Nets” paper [17] extends the traditional GCNN by employing multiple message aggregation functions. The operator first computes several independent aggregations of the messages using their mean, maximum, minimum and standard deviation, then additionally scales these aggregations based on the degree of the message-receiver node:

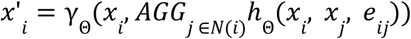

where *x_i_* is the embedding vector of the i-th node, *e_ij_* is the embedding of the edge between i-th and j-th nodes, N(*i*) returns the neighbors of i-th node, ℎ_Θ_ and γ_Θ_ are MLPs. *AGG* is the PNAConv aggregation applied to the output of ℎ_Θ_ and is defined as follows:

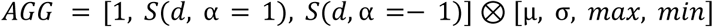

where ⊗ is the tensor product, d is the degree of the node, 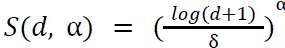 δ is a normalization parameter computed over the training set.

GENConv implements the Generalized Graph Convolution (GENConv) from the “DeeperGCN: All You Need to Train Deeper GCNs” [18] paper. It proposes using aggregation functions that unify the properties of multiple basic aggregations (for example, softmax to unify mean and max). In our work, we used softmax to aggregate the node messages:

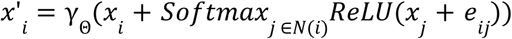

The notation of *x_i_*, *e_ij_*, N(*i*), γ_Θ_ is the same as in PNAConv.

TransformerConv is from “Masked Label Prediction: Unified Message Passing Model for Semi-Supervised Classification” paper [19]. It computes multi-head attention on neighbor node embeddings, aggregates the attention head outputs independently and uses their concatenation as the message:

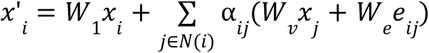

where *W_i_* is a weights matrix, and α*_ij_* are the attention coefficients obtained via attention (d is the size of embeddings *x_i_*):

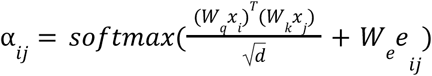

ResGatedGraphConv implements the residual gated graph convolutional operator from the “Residual Gated Graph ConvNets” paper [20]:

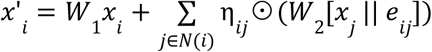

where || is the concatenation operator, ⊙ is the Hadamard pointwise multiplication operator, *W_i_* is a weights matrix, and 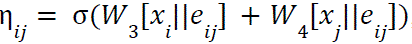, with σ denoting the sigmoid function.

#### Node and edge representations

We construct the input graph with the atoms as the nodes and the edges of the graph are defined based on the distance between atoms or the presence of a bond. If atoms *i* and *j* had distance below 4Å or had a chemical bond, we added an edge in the graph between the corresponding nodes.

In our ensemble models, each node feature is a trainable embedding for the corresponding atom type. We additionally tested pre-trained Uni-Mol node features [63], which also showed success in tasks that rely on molecular 3D properties. However, Uni-Mol employs parameter-heavy Transformer layers and significantly slowed down the inference time, while yielding very small or no gains for the selected models.

We encode each edge using the concatenation of 3 feature sets:

1. Embedding for each unique atom pair (e.g., “H-H”, “C-H”, or “O-H”).
2. Embedding for each edge type. We used 7 different edge types: 6 types indicating the bond (SINGLE, DOUBLE, TRIPLE, AROMATIC, IONIC, HYDROGEN), and another UNSPECIFIED type when the node atoms do not have a bond, but their inter-atom distance is lower than 4Å.
3. Expanded version of rotation-invariant Point Pair Features (PPF):
  a. distance between source and receiver nodes (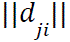, where *d*_*ji*_ denotes the difference vector between points)
  b. angle between *d*_*ji*_ and the surface normal vector of node i
  c. angle between *d*_*ji*_ and the surface normal vector of node j
  d. angle between the surface normal vectors of nodes i and j

We expanded the distance and angles to a 16-dimensional embedding, using 8-class one-hot encoding based on evenly spaced boundaries for each feature and a linear layer on top of each.

The performance of explored input features alternatives is provided in the following section.

#### Node and edge feature representation alternatives

For the selected 5 models we attempted to enrich their edge representations using PPF. The evaluation on the validation set demonstrated that these features consistently improved the MAE score (Table 4). Therefore, in all following experiments we used them as part of the models.

**Table 4.** Validation results for different edge representations.

| Model | Edge representation |  |  |  |
| --- | --- | --- | --- | --- |
|  | Edge type + Edge distance |  | Edge type + Edge distance + PPF |  |
|  | Valid RMSE | Valid MAE | Valid RMSE | Valid MAE |
| ResGatedGraphConv | 5.65 | 3.47 | **4.3** | **2.38** |
| GENConv | 5.11 | 2.96 | **3.65** | **1.87** |
| GeneralConv | 8.34 | 5.00 | **6.96** | **3.72** |
| TransformerConv | 5.46 | 3.38 | **5.01** | **2.24** |
| PNAConv | **4.37** | 2.85 | 4.81 | **2.23** |
| *Average* | *5.79* | *3.53* | *4.95* | *2.49* |

With the aim of finding more efficient feature representations, we additionally trained the models with Uni-Mol node features. We tested 3 feature configurations: (i) only trainable atom type embeddings, (ii) Uni-Mol features and trainable atom type embeddings, (iii) only Uni-Mol as node features. In both configurations, Uni-Mol embeddings were frozen during training. The addition of Uni-Mol embeddings did not improve ResGatedGraphConv, GENConv, TransformerConv models, and only showed slight MAE improvements for GeneralConv and PNAConv models (Table 5). During the training, we observed that Uni-Mol-based models started off relatively well on validation metrics, but then plateaued more quickly than the models without them (Figure 7). Thus, we decided not to use Uni-Mol in further experiments as it significantly increased the inference time and memory requirements without obvious improvement in validation metrics.

**Figure 7.**
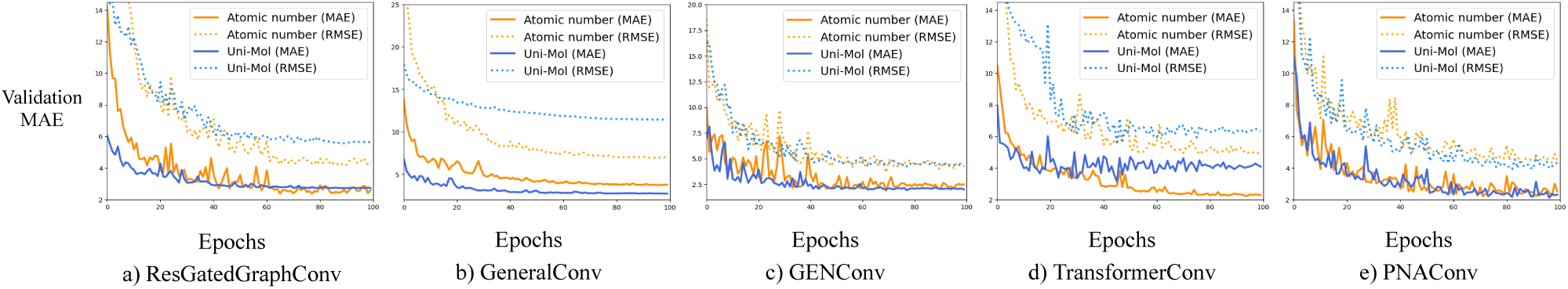
Impact of node feature selection on model training.

**Table 5.** Validation results for different node representations.

| Model | Node representation |  |  |  |  |  |
| --- | --- | --- | --- | --- | --- | --- |
|  | Atom type |  | Atom type + Uni-Mol |  | Uni-Mol |  |
|  | Valid RMSE | Valid MAE | Valid RMSE | Valid MAE | Valid RMSE | Valid MAE |
| ResGatedGraphConv | **4.3** | **2.38** | 5.64 | 2.73 | 6.04 | 2.73 |
| GENConv | **3.65** | **1.87** | 4.73 | **1.87** | 4.24 | 1.92 |
| GeneralConv | **6.96** | 3.72 | 8.77 | **2.62** | 11.46 | 2.72 |
| TransformerConv | **5.01** | **2.24** | 5.48 | 3.14 | 7.97 | 2.98 |
| PNAConv | 4.81 | 2.23 | **3.99** | **2.12** | 12.25 | 3.76 |
| *Average* | *4.95* | *2.49* | *5.72* | *2.50* | *8.39* | *2.82* |

PPF edge representations rely on the surface normal of the atoms as the anchor vector to calculate the angles. To avoid the relatively expensive computation of the normals, we additionally tested an alternative way of PPF calculation where the anchor vector for each atom is the normalized sum of the differences from its neighboring atoms’ position (Table 6). We refer to the original PPF as PPF-Normal hereafter, and the difference vector-based alternative is denoted as PPF-Diff. SDDF models based on each variant of PPF are respectively called SDDF-Normal and SDDF-Diff.

**Table 6.** Validation results for different PPF feature anchor vector calculation methods.

| Model | Edge representation |  |  |  |
| --- | --- | --- | --- | --- |
|  | PPF-Normal |  | PPF-Diff |  |
|  | Valid RMSE | Valid MAE | Valid RMSE | Valid MAE |
| ResGatedGraphConv | 4.3 | 2.38 | **2.70** | **1.87** |
| GENConv | 3.65 | 1.87 | **2.52** | **1.76** |
| GeneralConv | 6.96 | **3.72** | **5.63** | 3.80 |
| TransformerConv | **5.01** | **2.24** | 6.59 | 4.51 |
| PNAConv | 4.81 | 2.23 | **2.49** | **1.75** |
| *Average* | *4.95* | *2.49* | *3.99* | *2.74* |

The final models that were presented in the main manuscript use PPF-Diff version of the features.

#### Model training hyperparameters

For the initial model selection procedure, we trained each model for up to 100 epochs, using Mean Absolute Error loss function and Adam optimizer with 1e-4 initial learning rate. In the experiment for the evaluation of SDDF sampling strategies, we trained the models for up to 200 epochs. For the selected 5 architectures, we trained their final versions for up to 500 epochs. In each training run, the best checkpoint was selected based on the MAE score on the validation set. For regularization we use dropout with 0.15 probability. Mini-batch size is 128, where each sample is a graph for a single conformation. Unless specified otherwise, at the initial model selection stage we used the models’ default parameters and did not perform thorough hyperparameter tuning for each model separately.

We trained the models with targets represented in Hartree units. For all training runs, we shifted each molecule’s target energy by subtracting the estimated self-interaction atomic energies. The self-interaction atomic energies of each atom type were estimated via a linear regression model, where the atom type counts of a molecule are the input features and its total energy is the target. During the active learning simulation experiments, the estimated energies were re-calculated before every simulation round using the available Seed and SDDF sets. For the released energy prediction models that were benchmarked on our test set, we used the following self-interaction energies for each atom type:

H: –0.60210 Eh

C: –38.10081 Eh

N: –54.73759 Eh

O: –75.20245 Eh

S: –398.13982 Eh

Cl: –460.18514 Eh

Br: –2573.77264 Eh

F: –99.82343 Eh

While we trained the models using Hartree units, all evaluation results are reported in kcal x mol-1 unless specified otherwise.

#### Energy models’ performance relative to the state-of-the-art

We compared our individual conformational energy prediction models, as well as the ensembles, with the ANI-2x ensemble and the GemNet model trained on NablaDFT. Since ANI-2x was not trained on molecules containing bromine atoms, we present the evaluation results of our models on both the full test set and the subset without bromine-containing molecules (Table 7). Even without ensembling, the majority of the individual models outperformed ANI-2x on the test set.

**Table 7.** Comparison of the performance of ANI-2x and SDDF ensemble models on our test set (MAE in brackets indicates the results on the test subset excluding examples with Bromine atoms).

| Model |  | Test RMSE* | Test MAE* |
| --- | --- | --- | --- |
| SDDF-Normal | PNAConv | 4.14 (4.18) | 2.53 (2.52) |
|  | ResGatedGraphConv | 4.25 (4.26) | 2.77 (2.75) |
|  | GENConv | 4.23 (4.32) | 2.28 (2.27) |
|  | GeneralConv | 6.90 (6.89) | 4.11 (4.09) |
|  | TransformerConv | 5.08 (5.16) | 2.72 (2.71) |
|  | Ensemble {PNAConv, ResGatedGraphConv, GENConv} | 3.78 (3.83) | 2.17 (2.16) |
|  | Ensemble {all} | 4.13 (4.19) | 2.27 (2.26) |
| SDDF-Diff | PNAConv | 3.35 (3.38) | 2.06 (2.06) |
|  | ResGatedGraphConv | 3.39 (3.42) | 2.25 (2.24) |
|  | GENConv | 3.58 (3.60) | 2.25 (2.23) |
|  | GeneralConv | 6.02 (5.97) | 4.10 (4.08) |
|  | TransformerConv | 7.31 (7.34) | 5.12 (5.11) |
|  | Ensemble {PNAConv, ResGatedGraphConv, GENConv} | **2.89 (2.92)** | **1.83 (1.82)** |
|  | Ensemble {all} | 3.52 (3.54) | 2.31 (2.30) |
| ANI-2x** |  | — (5.07) | — (2.84) |

\*kcal x mol<sup>-1</sup>
\*\*Does not support Bromine

Since direct comparison of models trained on datasets with different DFT theory levels would not be valid (Figure 8), we decided to compare such models based on their ability to rank conformations using the predicted energy (Table 8). To keep the comparison fair, we did not use in this test any conformation generated by our methods. The SDDF ensemble trained on the SDDF train subset of the full dataset performed slightly better than the ANI-2x ensemble, but worse than NablaDFT in this ranking task. The lower score of the tested SDDF models can be attributed to the fact that their training set contained only 2-3 conformations per molecule on average, compared to 5+ for NablaDFT.

**Figure 8.**
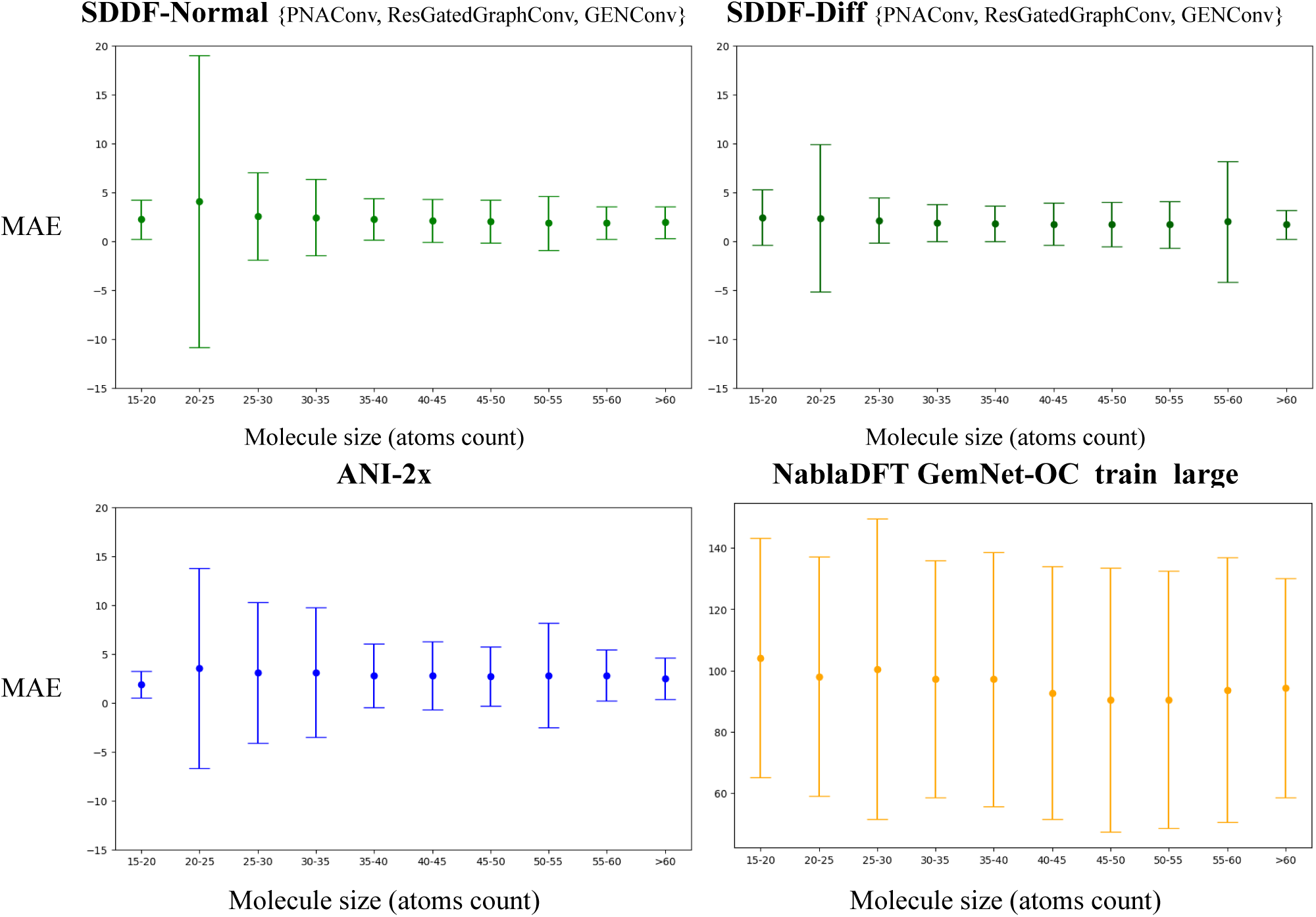
Comparison of models’ test set prediction errors based on molecule size.

**Table 8.** Comparison of the performance of ANI-2x, GemNet model trained on NablaDFT, and SDDF ensemble models on our test set (excluding examples with Bromine atoms and molecules with a single conformation) in the per-molecule conformation ranking task.

| Model | | $\rho_{\text{Spearman}}$ |
| --- | --- | --- |
| SDDF-Diff | PNAConv | 0.9029 |
|  | ResGatedGraphConv | 0.8920 |
|  | GENConv | 0.8996 |
|  | GeneralConv | 0.7662 |
|  | TransformerConv | 0.7312 |
|  | Ensemble {PNAConv, ResGatedGraphConv, GENConv} | 0.9177 |
| ANI-2x |  | 0.9120 |
| NablaDFT GemNet-OC train large |  | 0.9388 |

#### Evaluation of conformation sampling strategies

When evaluating the different sampling strategies, we independently ran the experiments with different random seeds. We performed the inference for each individual model, averaged the predictions among the ensemble models with the same random seed and calculated the validation MAE of these predictions. In Table 9, we report the average of these MAE results for the random seeds, and in Tables 10, 11, 12 we report the results for each random seed separately.

**Table 9.** Energy prediction test set MAE of the ML ensemble for different conformation sampling strategies.

| Sampling strategy | Seed | Step 1 | Step 2 | Step 3 | Step 4 |
| --- | --- | --- | --- | --- | --- |
| RANDOM | 5.72 | 6.20 | 4.51 | 4.52 | 4.25 |
| x5GENConv | 4.26 | 4.73 | 4.37 | 4.35 | 3.77 |
| SDDF-VAR | 5.72 | 5.21 | 4.08 | 4.00 | 3.87 |
| SDDF-LOSSFN | 5.72 | 4.10 | 3.97 | 3.89 | 3.73 |

**Table 10.**
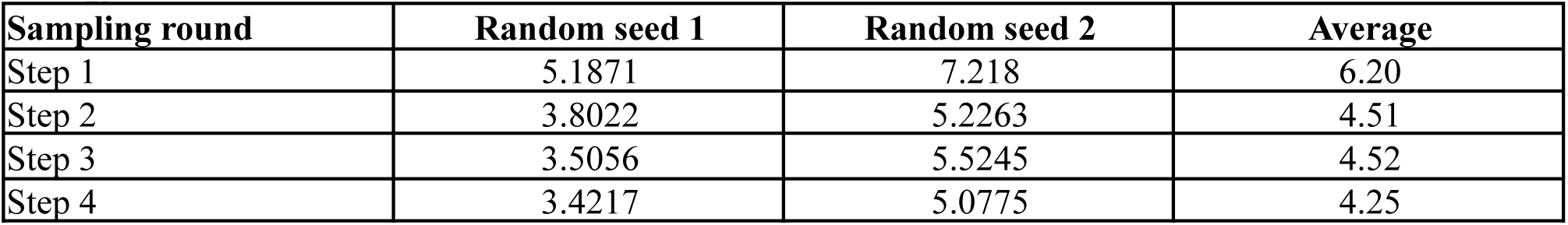
Energy prediction test set MAE of the ML ensemble for the random conformation sampling strategy.

**Table 11.**
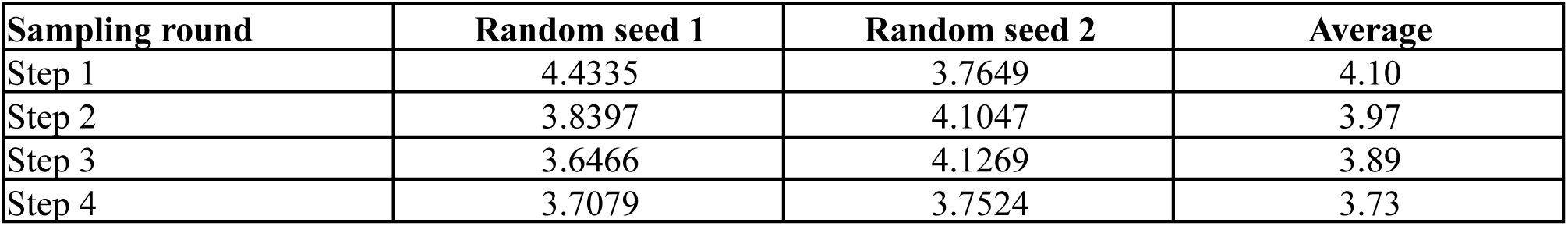
Energy prediction test set MAE of the ML ensemble for the model loss predictor-based conformation sampling strategy.

| Sampling round | Random seed 1 | Random seed 2 | Average |
| --- | --- | --- | --- |
| Step 1 | 4.4335 | 3.7649 | 4.10 |
| Step 2 | 3.8397 | 4.1047 | 3.97 |
| Step 3 | 3.6466 | 4.1269 | 3.89 |
| Step 4 | 3.7079 | 3.7524 | 3.73 |

**Table 12.** Energy prediction test set MAE of the ML ensemble for the variance-based conformation sampling strategy.

| Sampling round | Random seed 1 | Random seed 2 | Average |
| --- | --- | --- | --- |
| Step 1 | 4.1141 | 6.2959 | 5.21 |
| Step 2 | 3.8152 | 4.3525 | 4.08 |
| Step 3 | 4.0369 | 3.9711 | 4.00 |
| Step 4 | 3.9435 | 3.7882 | 3.87 |

#### Analysis of sampled molecules’ distributions

We compared the molecules selected by different sampling strategies in the active learning simulation setup. As illustrated in Figure 7, ensemble-based sampling algorithms do not demonstrate bias towards the selection of relatively large or small molecules and the selected conformations have similar molecule size distribution across all strategies. We also analyzed the distribution of the selected conformations’ energies and observed a similar absence of divergence between the different strategies’ distributions (Figure 8).

**Figure 9.**
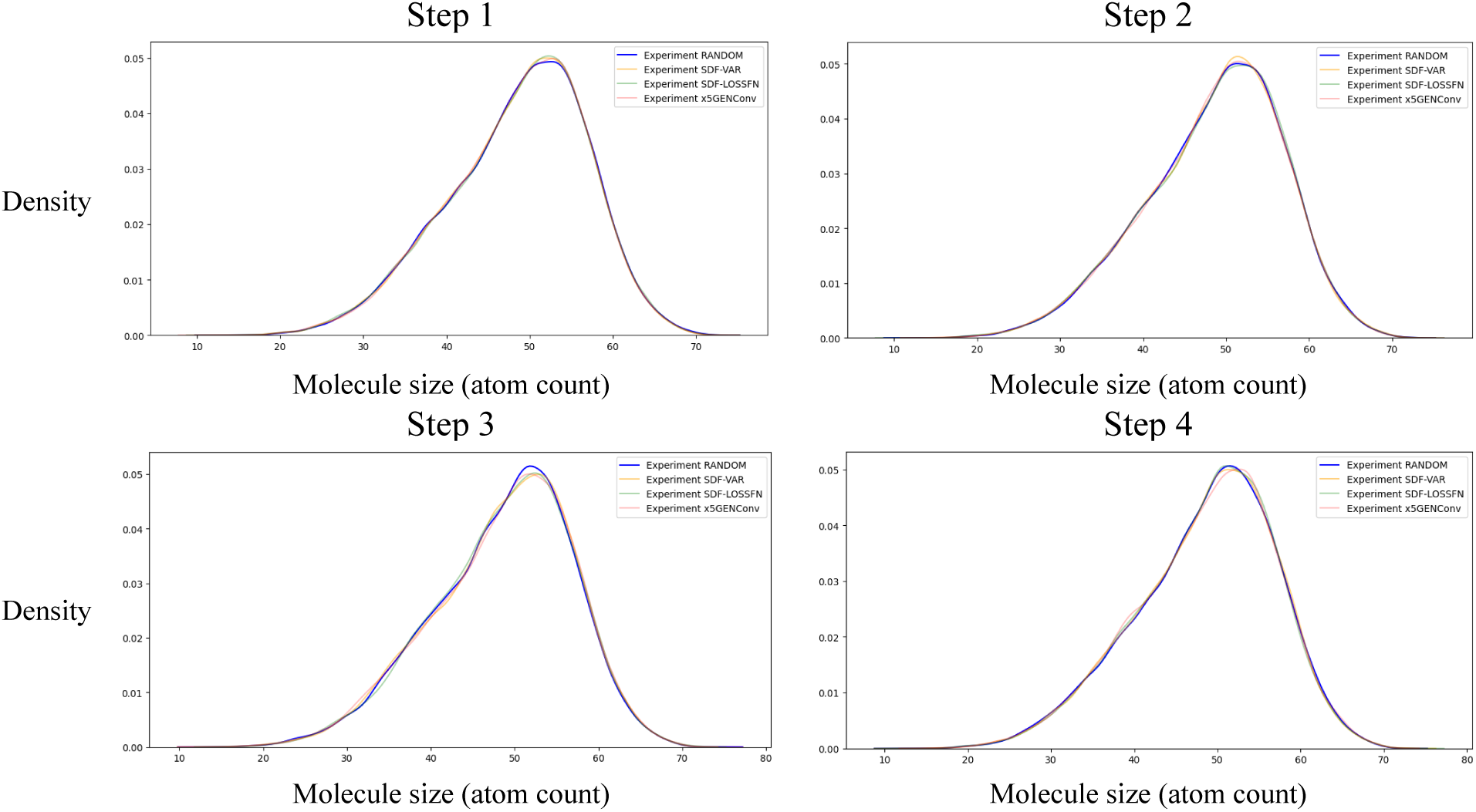
Comparison of molecule sizes for selected conformations at different steps of simulation for the 4 sampling strategies.

**Figure 10.**
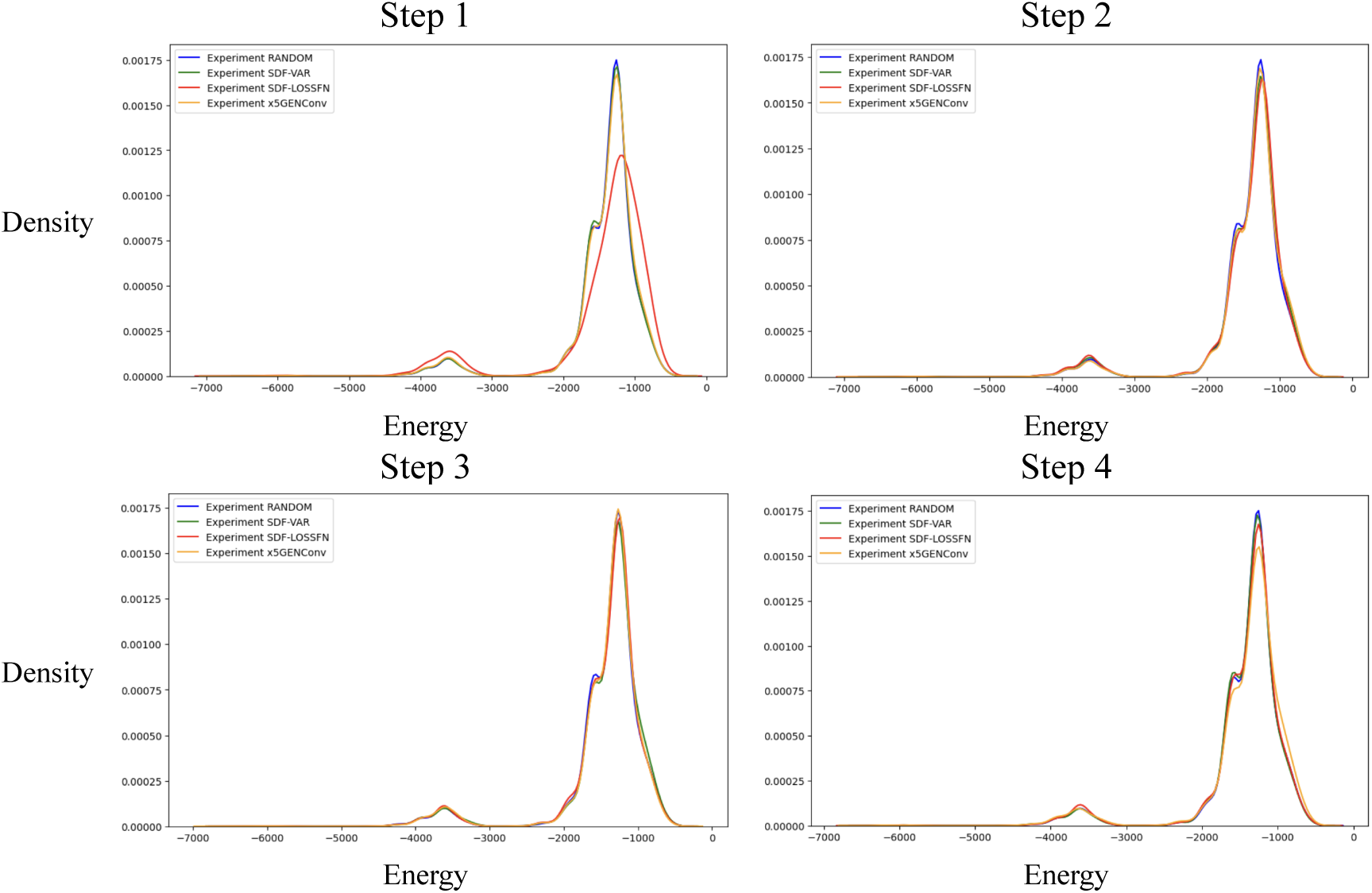
Comparison of energies for selected conformations at different steps of simulation for the 4 sampling strategies.

## Supplementary C. Energy prediction dataset and benchmark

The decision to create a new dataset for the energy prediction task is motivated by the limitations (particularly, the lack of diversity and the presence of train-test leakage) in the existing datasets, mentioned in the related work. As shown in Table 13, our SDDF dataset offers superior scaffold diversity compared to QM9, ANI-1, NablaDFT, and also better conformation diversity compared to QM9. We also believe it is very important to have a dataset based on molecules from a “real-world” database such as ENAMINE, which is one of the most popular compound libraries used in drug discovery projects. The dataset generation is ongoing (another very important factor) and we expect to continually increase the conformational diversity.

**Figure 11.**
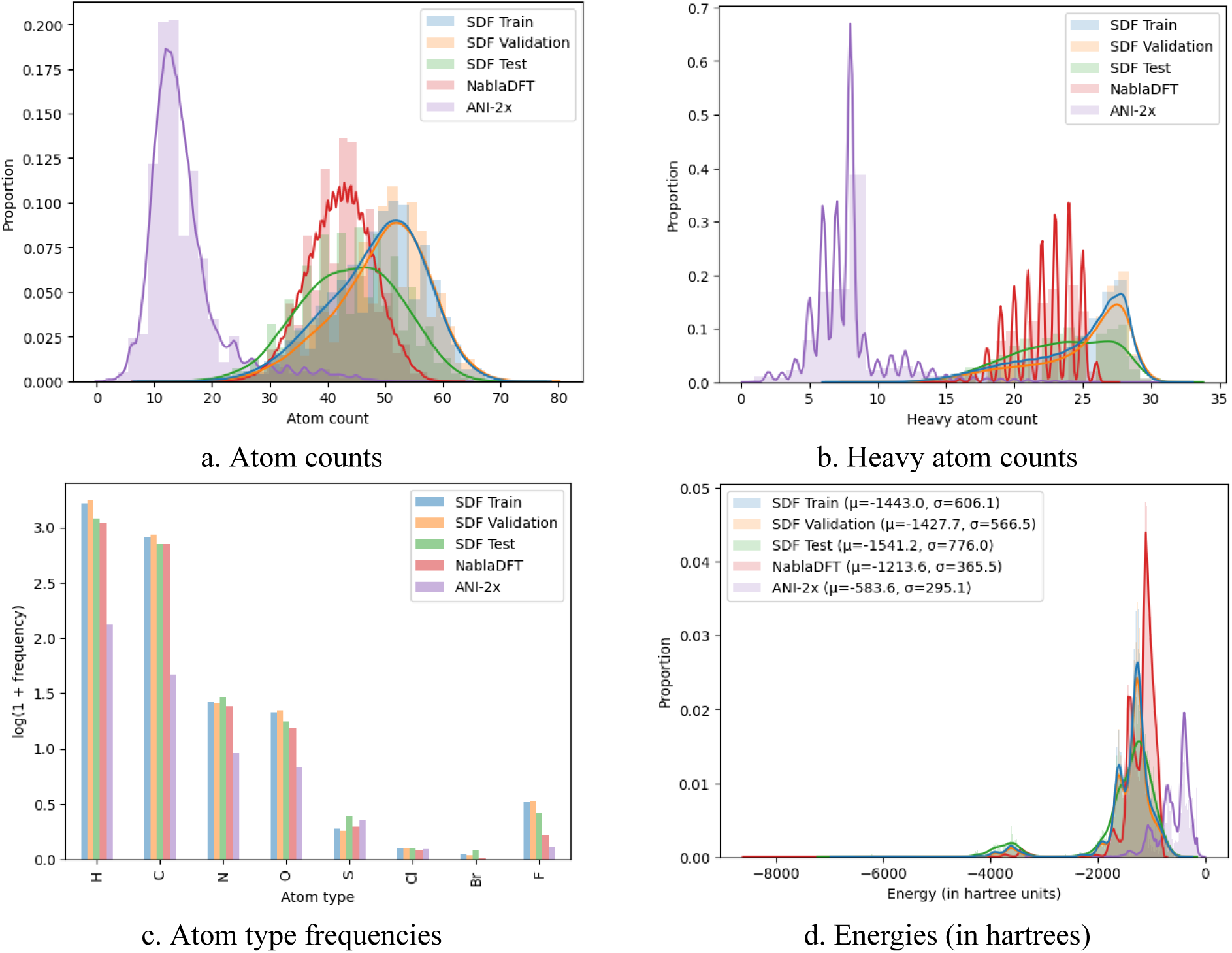
Comparison of atom count, heavy atom count and atom types distributions in ANI-2x, NablaDFT datasets and SDDF train, validation, test splits.

**Table 13.**
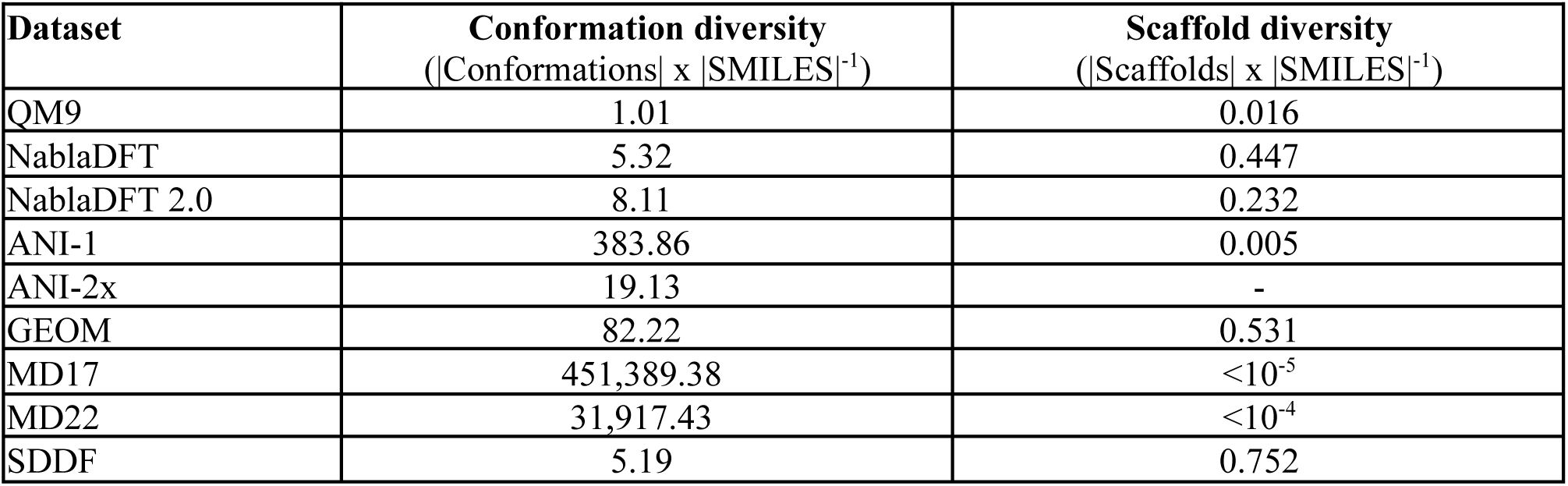
Diversity in molecular energy datasets. For ANI-2 [64], the scaffold diversity is not determined as the dataset does not provide bond information for its entries.

| Dataset | Conformation diversity<br>($\vert \text{Conformations}\vert \times \vert \text{SMILES}\vert ^{-1}$) | Scaffold diversity<br>($\vert \text{Scaffolds}\vert \times \vert \text{SMILES}\vert ^{-1}$) |
| --- | --- | --- |
| QM9 | 1.01 | 0.016 |
| NablaDFT | 5.32 | 0.447 |
| NablaDFT 2.0 | 8.11 | 0.232 |
| ANI-1 | 383.86 | 0.005 |
| ANI-2x | 19.13 | - |
| GEOM | 82.22 | 0.531 |
| MD17 | 451,389.38 | $<10^{-5}$ |
| MD22 | 31,917.43 | $<10^{-4}$ |
| SDDF | 5.19 | 0.752 |

Our full labeled dataset contains 2,170,553 conformations, including 535,338 that were generated using RDKit, another 1,151,936 that were generated using RDKit and optimized with MMFF94 force field, and 483,279 generated using MD on the RDKit conformations.

**Table 14.** Train, validation, and test datasets size statistics of our benchmark.

| **Dataset** | **Molecules** | **Scaffolds** | **Conformations** |
| --- | --- | --- | --- |
| Train | 274108 | 175179 | 638617 |
| Validation | 58293 | 38296 | 134732 |
| Test | 10063 | 8310 | 24890 |

